# BATF controls IFN-I production via DC-SCRIPT in plasmacytoid dendritic cells

**DOI:** 10.1101/2024.01.11.574638

**Authors:** Shafaqat Ali, Ritu Mann-Nüttel, Marcel Marson, Ben Leiser, Jasmina Hoffe, Regine J. Dress, Mahamudul Hasan Bhuyan, Patrick Petzsch, Karl Köhrer, Haifeng C. Xu, Philipp A. Lang, Shengbo Zhang, Michaël Chopin, Stephen L. Nutt, Judith Alferink, Stefanie Scheu

**Affiliations:** Institute of Immunology, Rostock University Medical Center, Rostock, Germany; Institute of Medical Microbiology and Hospital Hygiene, Medical Faculty and University Hospital Düsseldorf, Heinrich Heine University Düsseldorf; Cells in Motion Interfaculty Cluster, University of Münster, Münster, Germany; Department of Psychiatry, University of Münster, Münster, Germany; Division of Pulmonary Medicine, Department of Medicine, Faculty of Medicine & Dentistry, and Alberta Respiratory Centre, University of Alberta, Edmonton, Alberta, Canada; Institute of Systems Immunology, Hamburg Center for Translational Immunology (HCTI), University Medical Center Hamburg-Eppendorf, Hamburg, Germany; Department of Gynecology, University Medical Center Hamburg-Eppendorf, Hamburg, Germany; Biological and Medical Research Center (BMFZ), Medical Faculty and University Hospital Düsseldorf, Heinrich Heine University Düsseldorf; Department of Molecular Medicine II, Medical Faculty and University Hospital Düsseldorf, Heinrich Heine University Düsseldorf; Walter and Eliza Hall Institute of Medical Research, 1G Royal Parade, Parkville, VIC 3052, Australia; Department of Medical Biology, University of Melbourne, Parkville, VIC 3010, Australia; Department of Biochemistry, Monash Biomedicine Discovery Institute, Monash University, 15 Innovation Walk, Clayton, VIC 3800, Australia

**Author notes:** corresponding author: Stefanie Scheu **Email:**.

**Keywords:** plasmacytoid dendritic cells, type I interferon, BATF transcription factor, DC-SCRIPT, antiviral immunity

## Abstract

The basic leucine zipper ATF-like transcription factor (BATF) plays a pivotal role in coordinating various aspects of lymphoid cell biology, yet essential functions in plasmacytoid dendritic cells (pDCs) have not been reported. Here, we demonstrate that BATF deficiency leads to increased interferon (IFN) I production in Toll-like receptor 9 (TLR9)-activated pDCs, while BATF overexpression has an inhibitory effect. BATF-deficient mice exhibit elevated IFN-I serum levels early in lymphocytic choriomeningitis virus (LCMV) infection. Integrated transcriptomics and ChIP-seq approaches identified the transcriptional regulator DC-SCRIPT as a direct target of BATF that suppresses IFN-I promoter activity by interacting with interferon regulatory factor 7 (IRF7). Genome-wide association study (GWAS) analyses further implicate BATF in pDC-mediated human diseases. Our findings establish a novel negative feedback axis in IFN-I regulation in pDCs during antiviral immune responses orchestrated by BATF and DC-SCRIPT, with broader implications for pDC- and IFN-I-mediated autoimmunity.

**Significance Statement:** Type I interferons (IFN-I) are essential for antiviral immunity but, when dysregulated, can promote inflammatory and autoimmune diseases. Plasmacytoid dendritic cells (pDCs) are specialized producers of IFN-I, yet the mechanisms that prevent excessive IFN-I production remain incompletely understood. We identify the transcription factor BATF as a negative regulator of IFN-I production in pDCs and uncover a BATF-DC-SCRIPT regulatory axis that restrains this response. Following TLR activation, BATF directly induces DC-SCRIPT by binding its regulatory regions. DC-SCRIPT then interacts with the key IFN-I transcription factor IRF7 to limit IFN-I gene expression. Thus, BATF and DC-SCRIPT control a negative-feedback mechanism that modulates the magnitude and duration of pDC-mediated IFN-I responses, providing a framework for understanding and potentially therapeutically modulating IFN-I-driven inflammation.

## Introduction

Type I interferons (IFN-I) serve as a primary defense during viral infections by inducing a cell-intrinsic antiviral state and activating both innate and adaptive immunity (1). Plasmacytoid dendritic cells (pDCs) are recognized as producers of high amounts of IFN-I during viral infections and after activation of their key pattern recognition receptors (PRRs) Toll-like receptor (TLR) 7 and 9 (1–5). However, accumulating evidence suggests that IFN-I derived from pDCs can have both important protective and detrimental roles, particularly in the context of chronic viral infections such as HIV and autoimmunity (4, 6–8). Dysregulated pDC activation is also involved in promoting inflammatory and autoimmune diseases characterized by an IFN-I signature (2, 4, 6). Consequently, pDCs emerge as a significant target for immunotherapeutic interventions. It is therefore imperative to elucidate the hitherto poorly characterized molecular mechanisms governing the timing and magnitude of IFN-I responses.

BATF together with BATF2 and BATF3 comprise a subfamily within the larger group of AP-1 transcription factors (9). BATF plays an essential role in the development and function of different T cell subsets, including T helper (Th) type 2, Th9, Th17, follicular Th cells, CD8^+^ effector T cells and NKT cells (10–13). In addition, BATF influences the homeostasis and function of group 2 and group 3 innate lymphoid cells (ILCs), as well as class switch recombination in B cells (9, 14–16). However, no function of BATF in pDCs has been reported.

In this study, we show that basic leucine zipper transcription factor ATF-like (BATF) acts as a negative regulator of IFN-I production in pDCs by upregulating the transcription factor DC-SCRIPT. Using gain-of-function, loss-of-function and multiomics approaches, we demonstrate that BATF controls the expression of DC-SCRIPT in pDCs by binding to its cis-regulatory regions and remodeling its chromatin architecture. Our data indicate that DC-SCRIPT interacts with the transcription factor IRF7, thereby suppressing the IRF7-mediated induction of IFN-I. These findings significantly contribute to the understanding of the molecular mechanisms underlying the regulation of IFN-I expression in pDCs.

## Results

### TLR activation induces BATF expression in pDCs

To investigate BATF expression in pDCs, we analyzed its levels following in vivo TLR activation. CpG injection increased BATF expression in murine splenic pDCs, peaking at 12 h and remaining elevated for at least 24 h (Fig. 1a; Fig. S1a-c). In vitro, untreated bone marrow (BM)-derived pDCs showed minor amounts of BATF, which increased after TLR9 activation with CpG2216 (Fig. S1d, e) or 1668 (Fig. S1f, g). Flow-cytometry gating strategies for ex vivo (Fig. S2a) and in vitro (Fig. S2b) analysis of pDCs are shown in Fig. S2. Similar to TLR9 activation by CpG, TLR7 activation by the RNA virus LCMV also induced BATF expression in pDCs. In this setting, BATF expression peaked 96 h post infection (Fig. 1b; Fig. S3a), following the peak in serum type I IFN levels in infected mice (Fig. S3b).

**Fig. 1:**
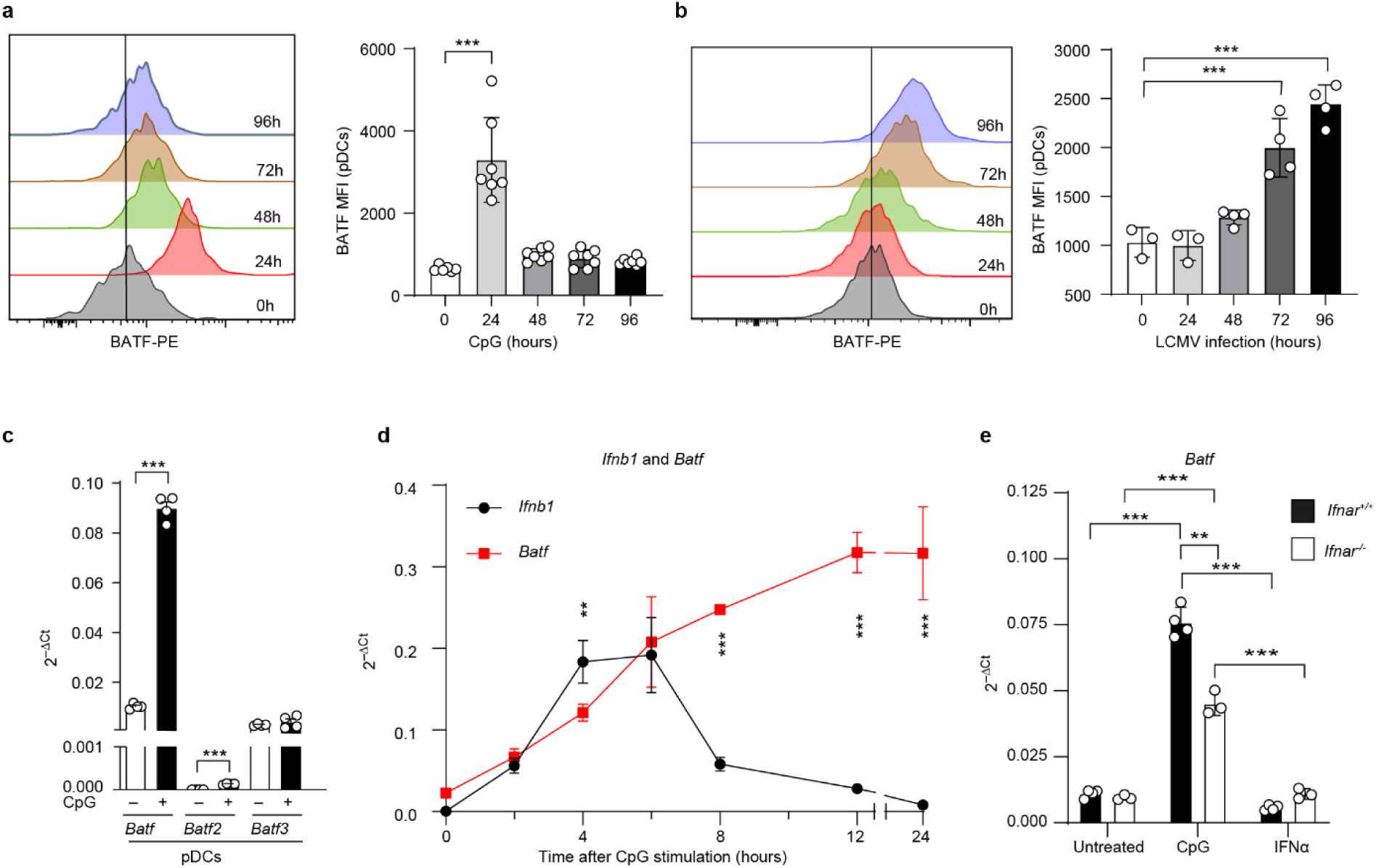
CpG stimulation and LCMV infection enhance BATF expression in pDCs. (**a, b**) Histograms (left) and bar charts (right) showing expression of BATF in splenic pDCs from C57BL/6 mice at steady state (0 h) or after in vivo CpG injection (**a**) or LCMV infection (**b**) for the indicated durations. The vertical grey line in the histograms indicates BATF MFI in resting pDCs. Bar graphs show the MFI (mean ± s.e.m) of BATF expression in splenic pDCs. (**c**) qRT-PCR analysis of relative *Batf*, *Batf2,* and *Batf3* mRNA expression in FACS-purified, BM-derived pDCs, resting or CpG-stimulated for 6 h. (**d**) qRT-PCR analysis of relative *Ifnb1* (black) and *Batf* (red) mRNA expression in FACS-purified, BM-derived, Flt3L-cultured pDCs at steady state (0 h) or after CpG stimulation for the indicated durations. (**e**) qRT-PCR analysis of relative *Batf* mRNA expression in FACS-purified BM-derived, Flt3L-cultured *Ifnar^-/-^* and WT (*Ifnar^+/+^*) pDCs at steady state or after 6 h of CpG2216 or IFNα4 stimulation. Data in (**c**-**e**) are means ± s.e.m., normalized to *Actb*. Statistical analyses: one-way ANOVA with Bonferroni’s multiple comparison test (**a**, **b**); two-tailed t test (**c**, **e**); multiple t test with Bonferroni correction (**d**). The data present pooled results from two experiments (**a**) or one representative experiment performed in triplicate or quadruplicate (n = 3-4 mice or independent cultures per group) from two (**a**-**c**, **e**) or three independent experiments (**d**) with comparable results. **: p < 0.01, ***: p < 0.001. MFI, median fluorescence intensity; pDCs, plasmacytoid dendritic cells; s.e.m., standard error of the mean.

Functional redundancy has been reported within the BATF family of transcription factors (17, 18). Therefore, we compared the expression levels of BATF, BATF2, and BATF3 in BM-derived pDCs. All three BATF factors are constitutively expressed in resting pDCs at low transcript levels, with *Batf2* only slightly above the detection threshold. *Batf* was the only transcript within the BATF family that increased markedly after TLR9 stimulation (Fig. 1c).

Analysis of transcriptome data (GSE68788) from IFNβ-producing pDCs purified from IFNβ^mob/mob^ reporter mice demonstrates a correlation between BATF expression and type I IFN production (19). Given this observed correlation, we investigated the time course of *Batf* expression in comparison to *Ifnb1* expression in these cells following CpG stimulation. The maximum of *Ifnb1* and multiple *Ifna* transcripts precedes the peak of *Batf* expression (Fig. 1d, Fig. S3c) in pDCs, suggesting that *Batf* expression could be affected by signals through the IFN-I receptor (IFNAR). To assess the impact of IFNAR signaling on *Batf* expression, we stimulated BM-derived pDCs from IFNAR-deficient mice and WT controls with CpG or recombinant IFNα2. CpG treatment increased *Batf* as well as *Ifnb1* mRNA expression in IFNAR-deficient pDCs, although to a lesser extent than in WT pDCs (Fig. 1e, Fig. S3d left panel). No direct induction of *Batf* transcripts by IFNα was found in WT cells, while upregulation of the IFN-inducible gene *Isg15* was evident (Fig. S3d right panel). These findings indicate that signals initiated by TLR9 directly induce the expression of BATF in pDCs, while IFNAR signaling alone does not play a major role in *Batf* induction but may contribute to the overall amounts of BATF in pDCs in the presence of TLR activation.

### BATF dampens IFN-I expression in pDCs

The delayed expression of BATF in comparison to IFN-I led us to hypothesize a modulatory role of BATF on IFN-I production rather than a role in its induction. Indeed, early after CpG stimulation, BM-pDCs from BATF-deficient (*Batf^-/-^*) mice expressed significantly more IFNβ and IFNα at the transcript (Fig. 2a) as well as protein (Fig. 2b) levels compared to *Batf^+/+^* controls. BATF deficiency also increased IL-12 and TNFα production in these cells, whereas other cytokines and chemokines were unchanged or reduced, including IL-18, IL-6, IL-10, eotaxin, MIP-1α, and MIP-1β (Fig. S4a).

**Fig. 2:**
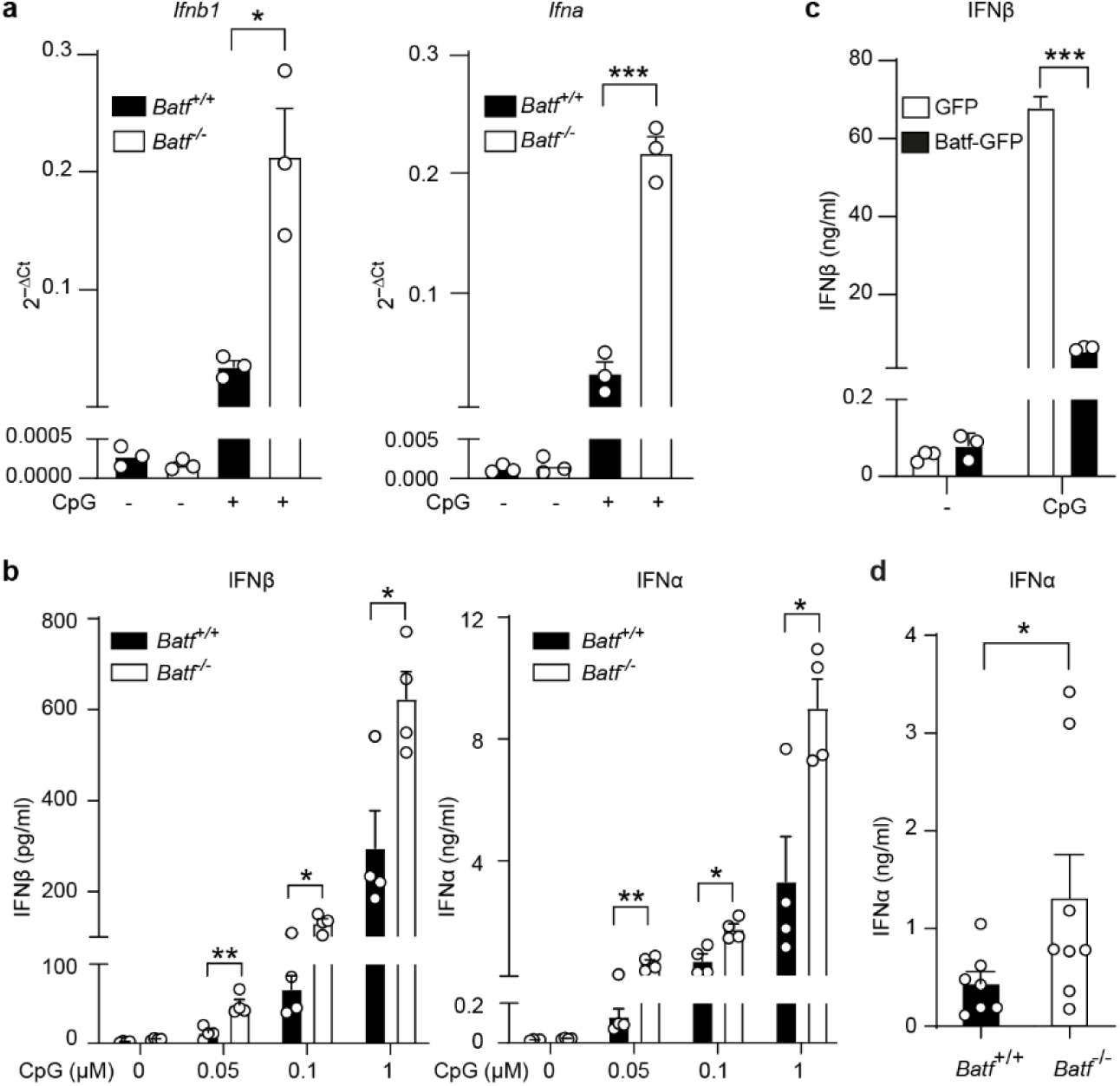
BATF negatively regulates IFN-I expression in pDCs. (**a**) qRT-PCR analysis of relative *Ifnb1* (left) and multiple *Ifna* (right) transcripts in untreated (-) or CpG-stimulated (2 h), FACS-purified BM-derived pDCs from WT (*Batf^+/+^*) and *Batf^-/-^* mice. (**b**) ELISA quantification of IFNβ (left) and IFNα (right) in supernatants of resting (0) and CpG-stimulated (indicated concentrations in µM; 4 h) FACS-purified, BM-derived pDCs. from *Batf^+/+^* and *Batf^-/-^* mice. (**c**) ELISA quantification of IFNβ in supernatants of FACS-purified murine Hoxb8-Fl pDCs overexpressing GFP or BATF-GFP, untreated (-) or CpG-stimulated for 16 h. (**d**) Serum IFNα levels 6 h after systemic application of CpG (2216) in *Batf^-/-^* (n = 8) and *Batf^+/+^* (n = 7) littermates. Data are means ± s.e.m (**a-d**) and normalized to *Actb* (**a**). Statistical analyses: unpaired two-tailed t test with Welch’s correction (**a**), unpaired multiple t tests using the Holm-Sidak correction (**b**), two-way ANOVA followed by Bonferroni’s multiple comparisons test (**c**), one-sided Mann-Whitney U test (**d**). Data represent one of three independent experiment performed using 4 independent cultures per group (**a**, **b**) or pooled data from three independent experiments (**c**, **d**). *: p < 0.05, ***: p < 0.001. BM, bone marrow; s.e.m, standard error of the mean.

Corroborating our findings in BATF-deficient pDCs, overexpression of BATF in Hoxb8-Fl-derived pDCs resulted in diminished IFNβ production compared to controls after CpG stimulation (Fig. 2c). Additionally, higher levels of IFNα were detected in the serum of BATF deficient mice compared to *Batf^+/+^* littermates (Fig. 2d) after systemic CpG stimulation.

Using IFNβ^mob/mob^ reporter mice, we found increased frequencies of IFNβ-producing pDCs in vitro following CpG stimulation under BATF-deficient conditions (Fig. S4b, c). In vivo, their frequency was unchanged, but their absolute number increased after CpG injection (Fig. S4d, e), consistent with the elevated IFN-I response. Collectively, our data suggest that both in vitro and in vivo, higher absolute numbers of IFNβ-producing pDCs contribute to the increased type I IFN levels observed in BATF deficiency.

In summary, loss- and gain-of-function experiments defined BATF as a negative regulator of IFN-I expression in pDCs. These data from both in vitro and in vivo experiments define the modulatory nature of BATF in early systemic virus infection-like conditions.

### BATF modulates the IFN gene signature in pDCs

To define the transcriptional effects of BATF deficiency, we performed RNA-seq on sorted WT and *Batf*^-/-^ BM-derived pDCs at steady state and 2, 6, and 12 h after CpG stimulation. We found 918 differentially expressed genes (DEGs) in unstimulated pDCs and 339, 1,489, and 1,570 DEGs at 2, 6, or 12 h of CpG stimulation, respectively, between the two genotypes (Fig. 3a). Pathway analysis showed enrichment of IFNα/β and cytokine signaling in resting BATF-deficient pDCs, with additional enrichment of IFN-I induction, IRF7 activation, and TLR-related pathways after CpG stimulation (Fig. 3b). GSEA confirmed upregulation of IFNα-response genes (Fig. 3c).

**Fig. 3:**
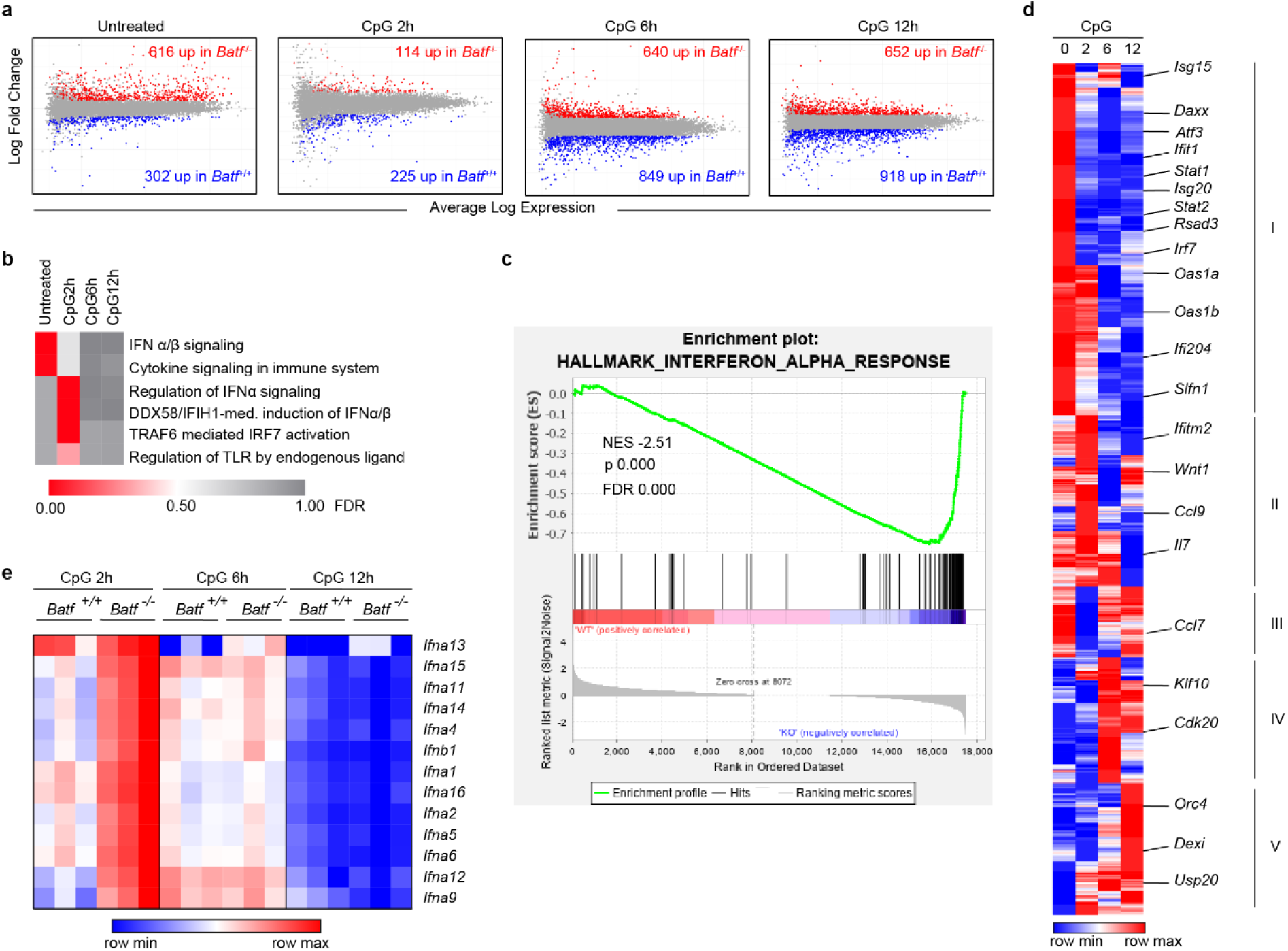
Activation of IFN-I signature genes and pathways in the absence of BATF. (**a**) MA plots showing global gene expression in murine BM-derived, Flt3L-cultured FACS-sorted *Batf*^+/+^ *and Batf*^-/-^ pDCs at steady state and after 2, 6, and 12 h of CpG stimulation. Differentially expressed genes (fold change ≥ |2|; FDR ≤ 0.05) are marked in red or blue. (**b**) Reactome pathway analysis of differentially expressed genes as in (**a**) with significance of the pathway (FDR) shown as a heatmap. (**c**) Gene set enrichment analysis (GSEA) of hallmark pathways using normalized RNA-seq expression data from untreated *Batf^+/+^* vs *Batf*^-/-^ pDCs and the gene set as permutation type. (**d**, **e**) Heatmap clustering of fold changes (**d**) and normalized expression values (**e**) for BATF-dependent IFN-I-stimulated genes (**d**) and IFN-I genes (**e**) in sorted *Batf^+/+^* and *Batf^-/-^* pDCs at steady state and after 2, 6, and 12 h of CpG stimulation.

More than half of the DEGs (504 out of 919) between untreated WT and BATF-deficient pDCs were annotated in the Interferome database v2.01 (20) as IFN-I-regulated genes (IRGs) in mice. Among them, 68% were upregulated in resting BATF-deficient pDCs (Table S1, Untreated). Similarly, at 2 h after CpG stimulation, 54% of the DEGs were IRGs (Table S1, 2 h CpG), while at later timepoints of stimulation, 6 h and 12 h, 42% qualified as IRGs (Table S1). Annotated IRGs (Table S1) were plotted in a clustered heatmap (Fig. 3d, selected genes are highlighted). Four of the five clusters show predominant upregulation under specific stimulatory conditions. Cluster I exhibited IRGs upregulated in untreated BATF-deficient pDCs, while clusters II, IV, and V represented IRGs differentially upregulated primarily after 2 h, 6 h, or 12 h of CpG stimulation, respectively. Cluster III comprised a small set of interferon stimulated genes (ISGs) showing upregulation under at least two stimulation conditions (Fig. 3d, Table S1). Interestingly, cluster I includes IRGs with tonic type I IFN-dependent expression. The majority of the genes in this cluster (e.g., *Daxx*, *Isg15*, *Isg20*, *Ifit1*, *Rsad2*, *Oas1a*, *Oas1b*, *Ifi204*, and *Slfn1*) have been reported to be less expressed in IFNAR-deficient B cells and macrophages (21), pointing to an increased secretion of tonic IFN-I from BATF-deficient pDCs. In addition, certain IRGs, designated as antiviral effectors (members of the *Gbp, Ifit*, *Mx*, or *Oas* gene family) and key regulators of IFN-I responses (*Irf7*, *Tlr7*, *Usp18*, etc.) were differentially upregulated in BATF-deficient pDCs in the absence of stimulation. Clusters II and III exhibit enrichment related to TLRs associated immune responses, ISGs linked to antigen presentation (e.g., *H2-Q4*, *H2-Q4*, *H2-Q7*, *H2-K1*) and proinflammatory functions (*Ccl7*, *Ccl9*, *Il7, Ccr2,* etc.), among others. Clusters IV and V encompass ISGs associated with various processes, such as metabolism (*Klf10*), cell cycle (*Orc4*), inflammation (*Dexi)*, and regulators of IFN-I responses (*Usp20*); however, genes in these clusters were not significantly enriched for associations with a specific function or process. BATF deficiency did not substantially alter the expression of most TLR9- or IFNAR-signaling components. However, *Irf7*, *Atf3*, *Stat1*, and *Stat2* were upregulated in resting pDCs (Table S2).

In line with our previous findings (Fig. 2), we found that pDCs stimulated with CpG showed significantly higher expression of all IFN-I genes, except for *Ifna13* at 2 h, but not at the later time points of 6 h and 12 h (Fig. 3e). Taken together, these data demonstrate that BATF significantly influences the global gene expression profile of both quiescent and activated pDCs. The most affected transcripts are IFN-I and other mRNAs linked to the IFN-I signature.

### BATF binds to the promoter of *Zfp366* and controls its expression in pDCs

To determine whether BATF regulates IFN-I genes directly or through other transcriptional regulators, we integrated ChIP-seq, ATAC-seq, and RNA-seq data from murine BM-derived pDCs.

BATF ChIP-seq in untreated pDCs from C57BL/6 mice and those stimulated for 2 h with CpG identified enrichment of the AP-1 and AICE1 motifs (Fig. S5a-f). Notably, both motifs were enriched to a higher extent after CpG stimulation of pDCs (Fig. S5a, right, c-f). Comparing the specific genomic locations with BATF binding, we found that BATF is mostly binding in similar proximal promoter and intron regions in untreated as well as 2 h CpG stimulated pDCs (Fig. S5b).

Integrated ChIP-seq and ATAC-seq analyses showed no BATF binding or significant changes in chromatin accessibility at most IFN-I gene loci, with the exception of *Ifna13* and *Ifna14* (Fig. 4a; Fig. S6a, b). Consistently, BATF overexpression did not alter *Ifnb1* or *Ifna4* promoter activity (Fig. S6c-g), despite a clear dose-dependent BATF expression was achieved in the transfected cells (Fig. S6g). Instead, after CpG stimulation, BATF did bind to several IFN-I response genes, such as *Isg15* and *Isg20* (Fig. 4a, Table S3). Taken together, these results negate direct binding and point to indirect suppression of IFN-I genes by BATF. Instead, BATF directly regulates the expression of a number of key transcription factors involved in DC development and function, including *Irf7*, *Irf8*, *Runx2*, *Spi1, Zfp366,* and *Spib* (Table S4, 5, 6).

**Fig. 4:**
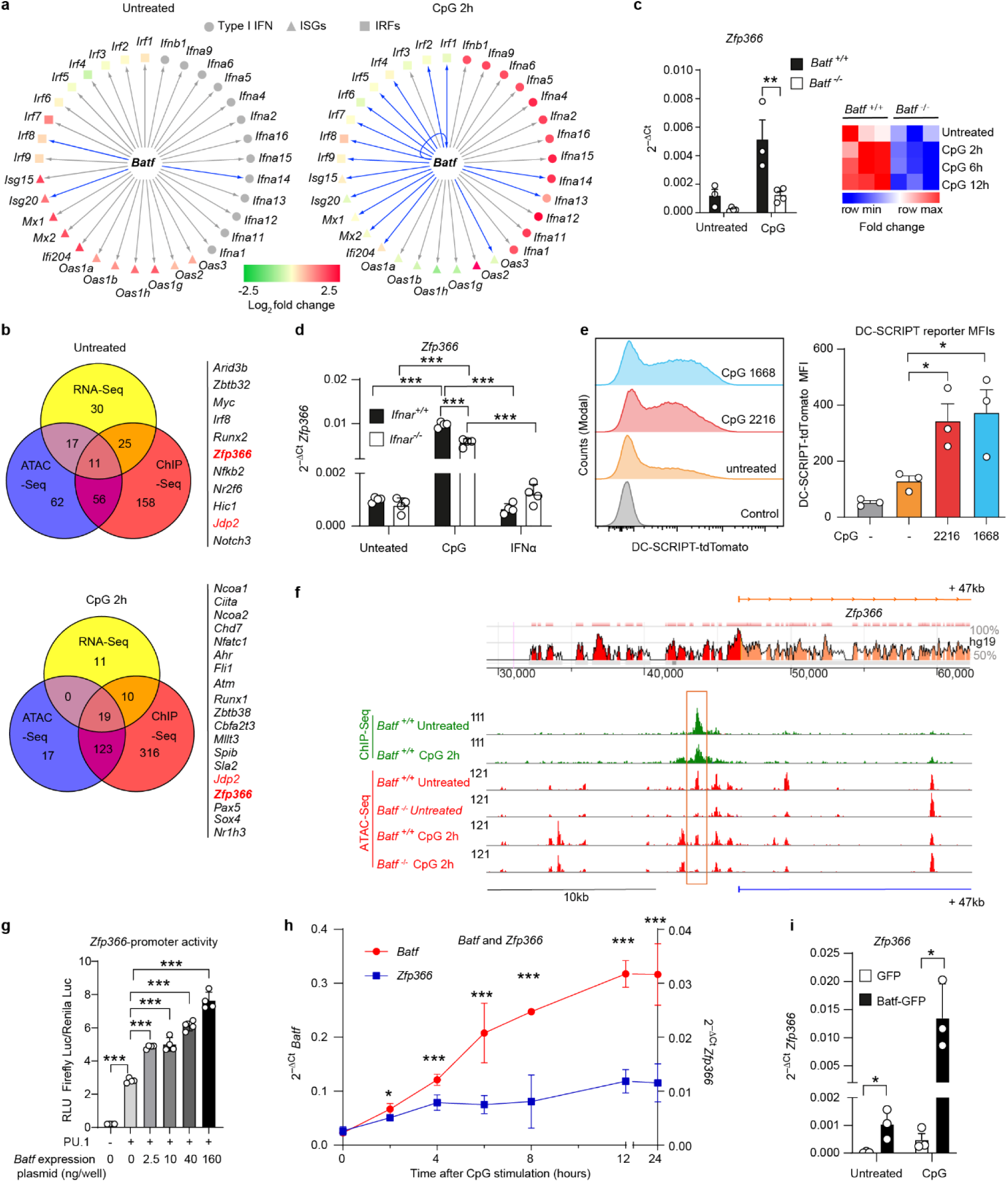
BATF binds to the *Zfp366* promoter and induces its expression in pDCs. (**a**) Cytoscape visualization of integrated RNA-seq and ChIP-seq data from WT and *Batf*^-/-^ BM-derived pDCs, untreated (left) or CpG-stimulated for 2 h (right). Shapes indicate gene groups (circles, IFN-I; triangles, ISGs; squares, IRFs); colors indicate expression changes in *Batf*^-/-^ versus WT pDCs (red, increased; green, decreased; grey, no expression). Arrow colors indicate BATF interaction (blue) or no interaction (grey), based on ChIP-seq. (**b**) Integrated RNA-seq, ChIP-seq and ATAC-seq data for mouse transcription factors in untreated (left) and 2 h CpG-stimulated (right) FACS-sorted murine BM-derived Flt3L-cultured pDCs visualized in Venn diagrams. Total number of genes among transcription factors is shown for differential expression of genes between WT and *Batf^-/-^* pDCs (RNA-seq, fold change ≥ |1.5| and FDR ≤ 0.05; EdgeR), direct BATF interaction with DNA (ChIP-seq peaks called after MACS) in WT pDCs and differential opening of chromatin between WT and *Batf^-/-^* pDCs (ATAC-seq, fold change ≥ |1.5| and FDR ≤ 0.05; DESEq2). Genes matching all three criteria are listed to the right of the Venn diagrams.(**c**) *Zfp366* expression measured by qRT-PCR in untreated and 6 h CpG-stimulated WT (*Batf*^+/+^) and *Batf*^-/-^ pDCs (left), and RNA-seq heatmap of normalized counts per million (CPM) in both genotypes at steady state and 2, 6 and 12 h after CpG stimulation (right). (**d**) *Zfp366* expression by qRT-PCR in untreated, CpG-stimulated or IFNα2-stimulated (6 h) WT (*Ifnar*^+/+^) and *Ifnar*^-/-^ pDCs. (**e**) Representative histograms (left) and MFI quantification (right) of *Zfp366*-tdTomato expression in untreated or 24 h CpG-stimulated BM pDCs from DC-SCRIPT reporter and WT control mice (grey bar). (**f**) ECR Browser view of the 5’ region of the mouse *Zfp366* gene (top; introns, pink; UTRs, yellow; conserved noncoding sequences, red) and IGV visualization of BATF ChIP-seq (green) and ATAC-seq (red) peaks in WT and *Batf*^-/-^ pDCs, untreated or CpG-stimulated for 2 h (bottom). Red boxes highlight BATF-binding and differential chromatin accessibility. (**g**) Effect of PU.1 and BATF on *Zfp366* promoter activity in dual-luciferase assays in transfected 293FT cells. Numbers below bars indicate BATF plasmid amounts (ng) per well. (**h**) Time-course qRT-PCR of *Batf* (red) and *Zfp366* (blue) expression in BM-derived pDCs, untreated (0) or CpG-stimulated for the indicated time span. (**i**) RT-qPCR showing the *Zfp366* expression in purified Hoxb8-FL pDCs overexpressing GFP or Batf-GFP, untreated or CpG-stimulated for 16 h.Data shown are from one representative experiment performed in duplicate (Chip-seq and ATAC-seq **a**, **b**, **f**), triplicate (**a**, **b** RNA-seq, **c**, **h**) or quadruplicate (**d**, **g**) out of three independent experiments with comparable results (**c**, **d**, **g**, **h**) or combined data from three independent experiments (**e**, **i**). Data in (**a**) are log_2_-fold changes; data in (**c**, **d**, **h**, **i**) are the mean ± s.e.m from 3-4 biological replicates normalized to *Actb*; data in (**g**) are means ± s.e.m firefly/Renilla luciferase ratios. Statistical tests: unpaired two-tailed *t*-tests (**c**, **e**, **i**), Bonferroni-corrected multiple *t*-tests (**d**), one-way ANOVA (**g**) or two-way ANOVA with Bonferroni-corrected multiple comparisons (**h**). *: p < 0.05, **: p < 0.01, ***: p < 0.001. s.e.m, standard error of the mean.

We hypothesized that transcription factors involved in BATF-mediated IFN-I suppression would likely fulfil the following criteria: (1) they exhibit BATF-dependent expression, (2) BATF binds to the regulatory region of the transcription factor in pDCs, and (3) the chromatin accessibility of the gene must be significantly altered by the loss of BATF in pDCs. Two genes, *Jdp2* and *Zfp366*, fulfilled the criteria in pDCs both at steady state and after TLR9 activation (Fig. 4b). We focused on *Zfp366*, the gene encoding the transcription factor DC-SCRIPT, because it is expressed in myeloid DCs, pDCs, and Langerhans cells (22). Importantly, no prior reports have documented its involvement in pDC biology or IFN-I production.

RT-PCR demonstrated reduced expression of *Zfp366* in BATF-deficient BM-derived pDCs compared to that in WT controls (Fig. 4c). Stimulation with recombinant IFNα did not influence the expression of *Zfp366* in *Ifnar^+/+^* or *Ifnar^-/-^* pDCs. Additionally, compared with those from WT cells, BM-derived pDCs from IFNAR-deficient mice exhibited comparable expression of *Zfp366* early (2 h) after CpG stimulation (Fig. S7). However, at later time points (6 h), we observed lower amounts of DC-SCRIPT-encoding transcripts in the absence of an IFNAR response (Fig. 4d). These data demonstrate that expression of *Zfp366* is largely independent of IFNAR responses in pDCs. However, in the presence of activated TLR9 signaling, IFNAR signaling may slightly contribute to *Zfp366* expression in pDCs. Corroborating these findings, we observed minor amounts of *Zfp366*-encoded DC-SCRIPT in pDCs at steady state in a DC-SCRIPT reporter mouse model. However, upon activation with CpG, DC-SCRIPT/tdTomato expression was elevated in pDCs (Fig. 4e).

Our ChIP-seq data revealed strong binding of BATF to the conserved noncoding sequence 1 (CNS1) of the *Zfp366* promoter (Fig. 4f, Fig. S8a), with major binding sites positioned at approximately -1270, -1400, -2500, and -3500 bp upstream of the transcription start site. Consistent with these findings, ATAC-seq analysis demonstrated greater chromatin accessibility at these BATF-bound regions in WT than in *Batf*^-/-^ pDCs (Fig. 4f). This suggests that regulation of chromatin structure in combination with direct binding of BATF to the *Zfp366* gene is responsible for BATF-mediated *Zfp366* expression.

Public data confirmed the binding of BATF family members to the *Zfp366* promoter sequences in T cell subsets and cDCs (Fig. S8b), with both public data and our ChIP-seq identifying BATF binding at regulatory regions RR2, RR3, and RR4 (Fig. S8b). We therefore generated *Zfp366* promoter-reporter constructs with deletions of regions containing BATF-binding motifs (Fig. S8c).

Data from the full-length *Zfp366* promoter (-3904 to +100 relative to the transcription start site) and its deletion mutants are presented in Fig. 4g and Fig. S8d, e. The *Zfp366* promoter contains multiple binding sites for ETS family transcription factors (e.g. PU.1 and Spi-B), which are constitutively expressed in pDCs. In HEK cells transfected with *Zfp366* reporter plasmids, we therefore overexpressed PU.1 along with BATF. As anticipated, PU.1 alone activated the *Zfp366* promoter to a certain extent (Fig. 4g). Addition of BATF expression further increased the promoter activity in a dose-dependent fashion (Fig. 4g), indicating direct involvement of BATF in regulating the DC-SCRIPT coding gene.

BATF overexpression increased *Zfp366* promoter activity, both in the absence (Fig. S8d) and presence (Fig. S8e) of PU.1, but only when the promoter retained RR1-4 (full-length) or RR1-3 (lacking RR4). Deletion of RR3 and 4 or RR2-4 abolished the additional effect of BATF, although PU.1 could still activate the promoter (Fig. S8d, e). These results demonstrate that BATF directly binds the *Zfp366* distal-promoter region and significantly contributes to its activation.

Together with ChIP-seq and *Zfp366* expression datasets from pDCs in the absence and overexpression of BATF, these reporter gene assay data provide independent and complementary evidence supporting BATF-mediated regulation of *Zfp366* expression in pDCs.

To further delineate the expression trajectory of DC-SCRIPT in activated pDCs, we analyzed the time course of *Batf* vs. *Zfp366* expression. We found that *Batf* transcript levels preceded *Zfp366* expression early after activation, supporting BATF-mediated expression of DC-SCRIPT in pDCs (Fig. 4h). In a gain-of-function situation, ectopic BATF expression in murine HoxB8-Fl pDCs significantly induced *Zfp366* expression even under steady state conditions, which was further enhanced upon CpG treatment (Fig. 4i). Taken together, our data point to BATF as a potent regulator of DC-SCRIPT expression in pDCs.

### DC-SCRIPT dampens IFN expression in pDCs

Next we characterized the role of DC-SCRIPT in IFN-I expression in pDCs. In *Ifnb1* and *Ifna4* reporter gene assays, overexpression of DC-SCRIPT strongly suppressed IRF7-induced *Ifnb1* (Fig. 5a) and *Ifna4* (Fig. 5b) promoter activity in a dose-dependent manner. The complementary approach of ectopic expression of DC-SCRIPT in Hoxb8-Fl pDCs led to reduced IFNβ production under both constitutive and CpG stimulation conditions (Fig. 5c), confirming the negative regulatory role of DC-SCRIPT in IFNβ expression. Consistent with previous studies (23, 24), DC-SCRIPT deficiency increased, whereas its overexpression decreased, pDC frequencies in in vitro cultures. Therefore pDC numbers and surface marker expression were normalized by FACS-purification across all experimental conditions. To exclude additionally developmental differences or cDC contamination in the sorted pDCs as explanations for altered IFN-I expression, we analyzed *Tcf4* and *Id2*, key pDC/cDC fate-determining transcription factors. Their mRNA ratios were comparable between DC-SCRIPT-overexpressing and control pDCs (Fig. S9a), indicating comparable developmental status. Although DC-SCRIPT regulates *Il12b* (IL-12p40) in cDC1s (24), its overexpression did not affect IL-12p40 secretion by purified pDCs (Fig. S9b), supporting a cDC1-specific function as reported earlier (24)

**Fig. 5:**
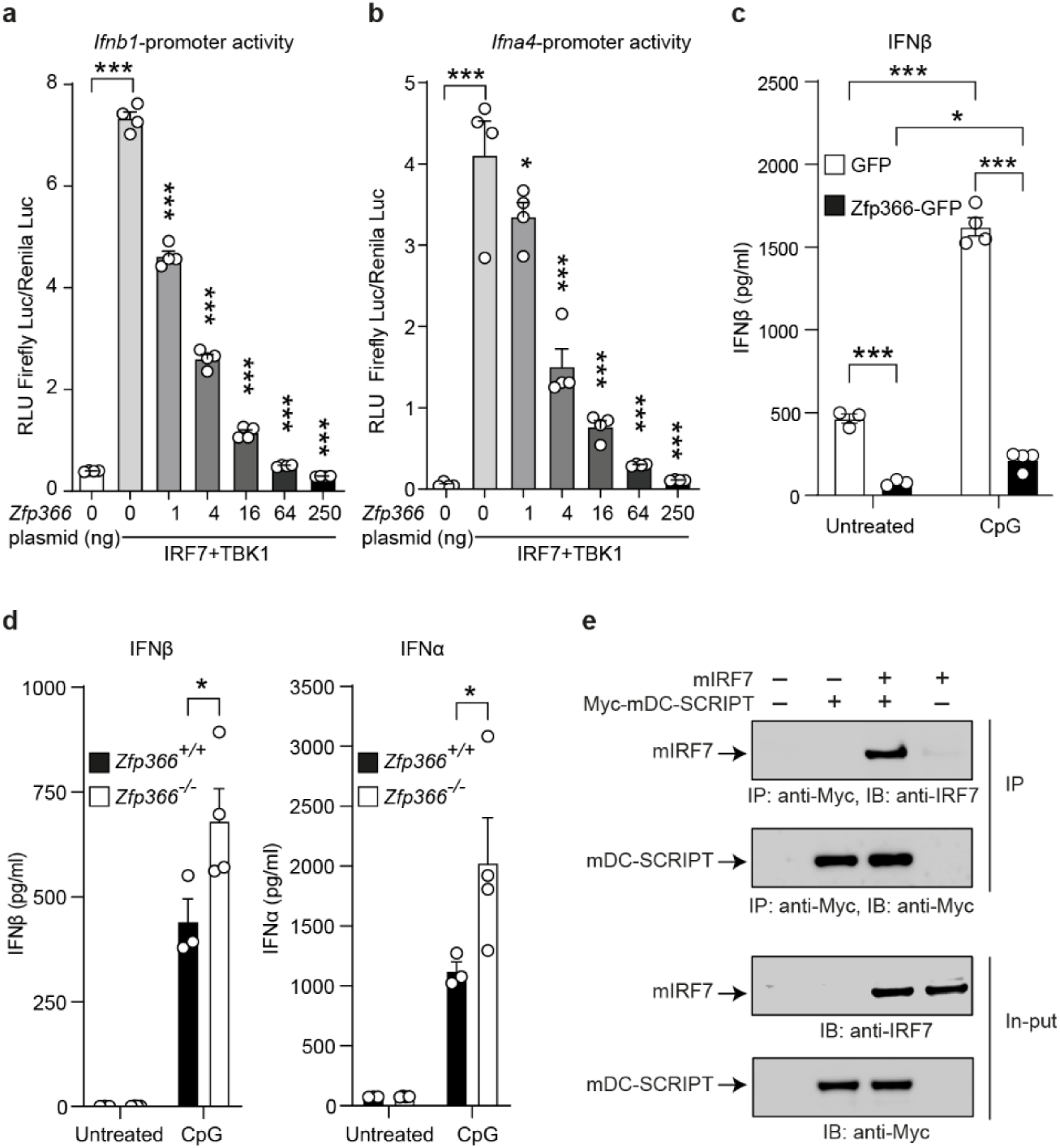
DC-SCRIPT interacts with IRF7 to dampen IFN-I expression. (**a, b**) Dual-luciferase reporter assays in transiently transfected 293FT cells showing the effect of DC-SCRIPT on IRF7-induced *Ifnb1* (**a**) and *Ifna4* (**b**) promoter activity. Numbers below bars indicate the amount of DC-SCRIPT plasmid (ng) per well. (**c**) IFNβ expression in FACS-purified murine Hoxb8-FL pDCs expressing GFP or DC-SCRIPT-GFP (Zfp366-GFP), untreated or CpG-stimulated for 16 h. (**d**) IFNα and IFNβ concentrations in supernatants of untreated or CpG-stimulated (12 h) FACS-purified, Flt3L-cultured pDCs from the BM of *Zfp366^-/-^*◊WT or WT◊WT BM chimeric mice measured by ELISA. (**e**) Co-immunoprecipitation of murine DC-SCRIPT and IRF7 in transfected 293FT cells using anti-Myc agarose beads and the indicated antibodies. Data in (**a**, **b**) are means ± s.e.m firefly/Renilla luciferase ratios. Data represent one of three (**a**-**c, e**) or two (**d**) independent experiments. Statistical significance was tested using one-way ANOVA with Fisheŕs LSD multiple comparisons test. *: p < 0.05, ***p < 0.001. s.e.m, standard error of the mean.

In keeping with the findings from gain-of-function experiments, we observed enhanced production of IFNβ and IFNα by CpG-stimulated BM-derived DC-SCRIPT-deficient pDCs compared with WT controls (Fig. 5d).

IRF7 in pDCs is indispensable for the rapid and robust IFN-I production upon pathogen recognition (25). Given that DC-SCRIPT reversed IRF7-induced *Ifnb1* and *Ifna4* promoter activation (Fig. 5a, b), we speculated that DC-SCRIPT functionally interacts with IRF7. Indeed, co-immunoprecipitation of murine IRF7 together with murine DC-SCRIPT supported the possible mechanism of DC-SCRIPT-mediated inhibition of IRF7 functions (Fig. 5e).

Together, these results support a model in which BATF-induced DC-SCRIPT interacts with IRF7 and limits IFN-I production in pDCs.

### Human DC-SCRIPT interacts with human IRF7

By analyzing *BATF* expression in previously published transcriptomics data (GSE93679)(26) for human pDCs (Fig. 6a), we found that *BATF* was upregulated along with IFN-I in human pDCs stimulated with CpG for 24 h. These data indicate a conserved expression pattern of mouse and human BATF in pDCs across the species.

**Fig. 6.**
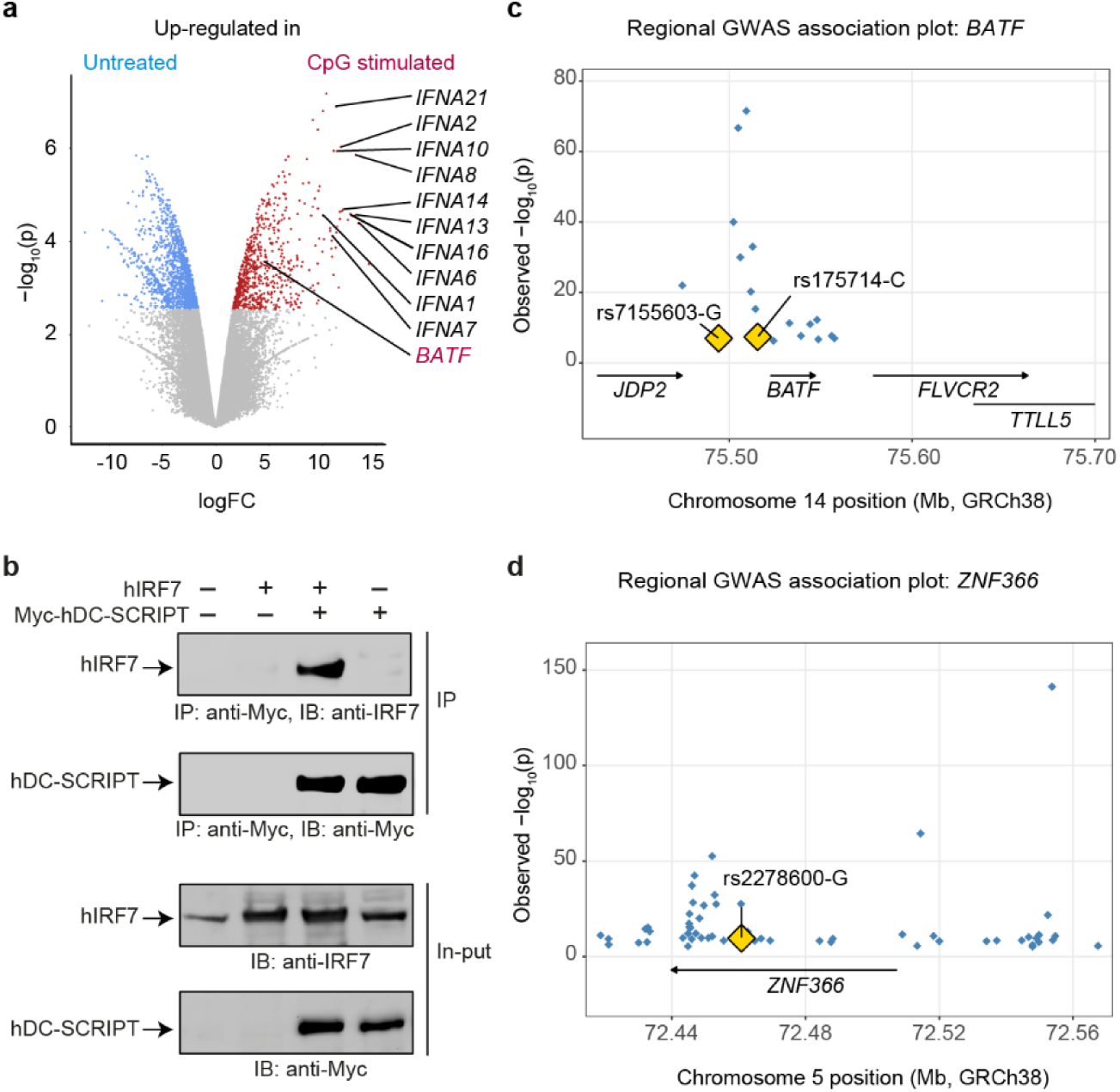
TLR9 activation induces BATF in human pDCs. (**a**) Volcano plot of gene expression in untreated and CpG-stimulated (24 h) human peripheral blood pDCs (GSE93679), reanalyzed using Biobase, GEOquery and limma. (**b**) Co-immunoprecipitation of human IRF7 and DC-SCRIPT, performed as in Fig. 5e. The data is representative of three independent experiments. (**c, d**) Regional plots showing GWAS Catalog associations (GRCh38) at the *BATF* (**c**) and *ZNF366* (**d**) loci. Each diamond represents a reported association (any trait), with the most significant entry per variant shown. Rheumatoid arthritis-associated variants rs7155603–G and rs175714–C upstream of *BATF* (**c**) and the missense variant rs2278600–G within *ZNF366* (***d***) are highlighted in yellow,. Arrows indicate gene orientation. TTLL5 extends beyond the plotted region (**c**).

DC-SCRIPT expression in humans has been reported in pDCs as well (27). Interestingly, all known functional domains are evolutionarily well conserved in murine and human DC-SCRIPT (28), as are the functional domains of IRF7 (29). Indeed, human DC-SCRIPT also co-immunoprecipitated with human IRF7 (Fig. 6b), supporting conservation of this protein interaction between mouse and human.

To unravel the potential genetic association of BATF and DC-SCRIPT with human health and disease, particularly in the context of IFN-I-mediated pathologies, we mined the publicly available GWAS Catalog (30). Our analysis identified two sequence variants, rs7155603-G and rs175714-C, located upstream of the human *Batf* gene (Fig. 6c, Table S7), and a missense variant within *ZNF366*, rs2278600-G (Fig. 6d, Table S7), associated with increased rheumatoid arthritis risk (Table S7), a disease in which IFN-I is discussed as a pathology driving entity (31).

Together, these findings demonstrate that *BATF* expression is induced by CpG stimulation in human pDCs and that human DC-SCRIPT interacts with IRF7.

### BATF dampens IFN-I expression during systemic viral infection

Next, we investigated the role of BATF early in viral infection in vivo, during which pDCs significantly contribute to IFN-I production (32). For this purpose, we infected WT and BATF-deficient mice with LCMV. Serum IFNα levels were elevated in *Batf^-/-^* mice on days 3 and 4 after LCMV injection (Fig. 7a; Fig. S10a), whereas IFNβ concentrations were significantly increased at day 4 (Fig. 7b; Fig. S10b), indicating a modulatory role of BATF in innate antiviral immune responses. Despite elevated IFN-I levels, splenic viral titers were comparable between genotypes (Fig. S10c), potentially reflecting impaired innate lymphoid cell (ILC) and T cell responses previously reported in BATF-deficient mice (12, 14). No detectable virus was found in the lungs, liver, or kidneys of either genotype at day 4 post-infection, possibly due to the low viral dose (data not shown).

**Fig. 7:**
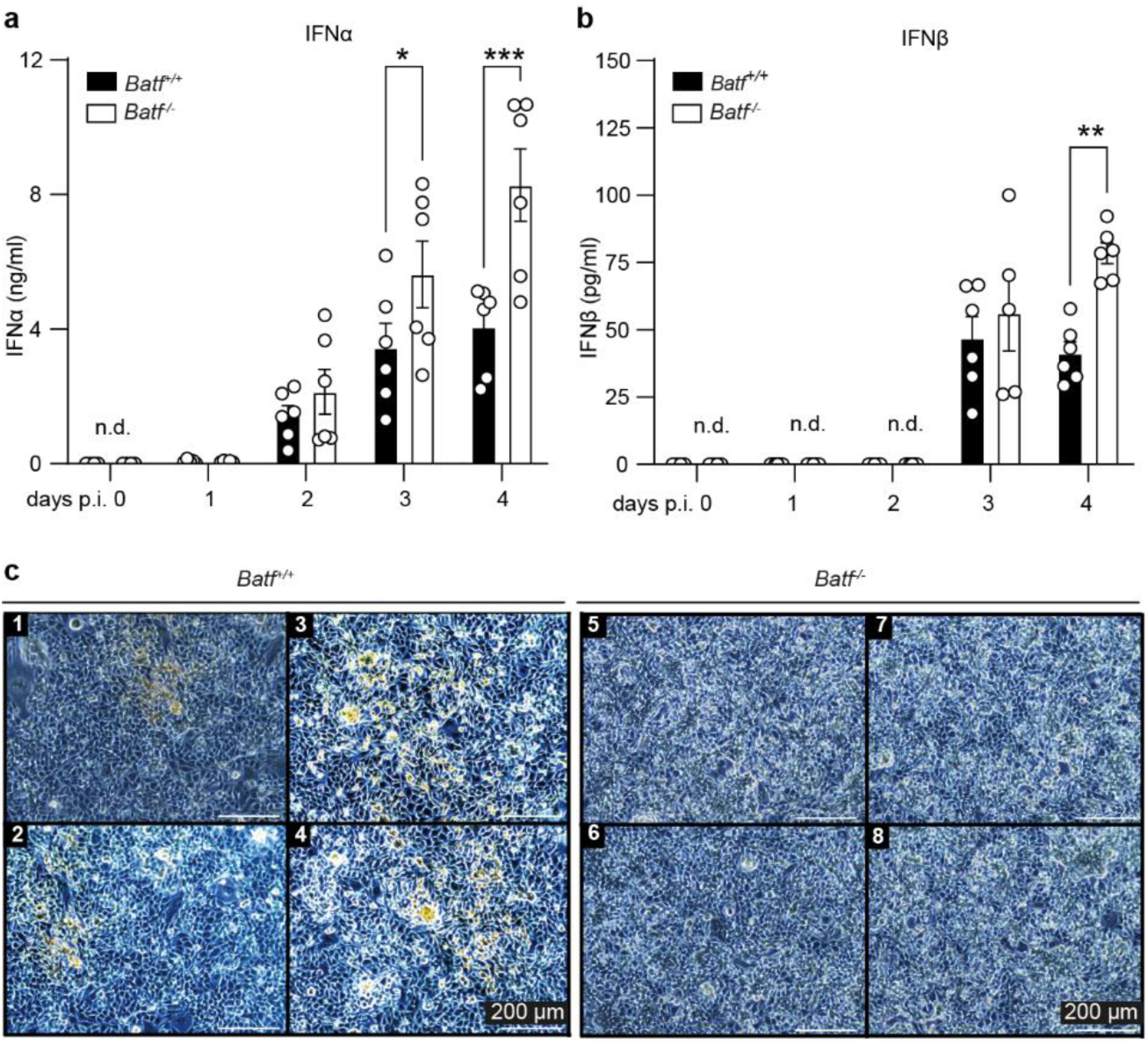
BATF dampens IFNα production during systemic LCMV infection. (**a**, **b**) *Batf^-/-^* mice and WT (*Batf^+/^*^+^) littermates were i.v. infected with 200 PFU of the LCMV-WE strain. Serum IFNα (**a**) and IFNβ (**b**) were measured by ELISA on days 1-4 post-infection. No IFN was detectable in uninfected mice, and IFNβ remained undetectable on days 1-2. (**c**) Representative images of MC57 fibroblasts cocultured with *Batf^+/+^* (1-4) or *Batf^-/-^* (5-8) FACS-sorted BM-derived Flt3L-cultured pDCs and infected with LCMV-WE. Data shown in (**a**, **b**) are means ± s.e.m. Statistical differences were analyzed using two-way ANOVA followed by Bonferroni’s multiple comparisons test; values below the detection limit were excluded. Data are representative of two experiments (n = 6). n.d., not detectable; s.e.m, standard error of the mean.

While under steady-state conditions, serum IFN-I was undetectable (Fig. 7a, b), BATF-deficient unstimulated pDCs showed increased expression of several ISGs (Fig. 3d). Consistently, lymph node expression of *Isg15*, *Isg20*, and *Mx1* was significantly increased in *Batf*^-/-^ mice compared with WT controls (Fig. S10d). These findings suggest enhanced basal ISG expression and potentially elevated tonic IFN-I signalling in BATF-deficient mice.

To exclude contributions from dysregulated ILCs and T cells in complete *Batf*-deficient mice, we generated pDC-specific *Batf* knockout mice by crossing *Batf*-floxed mice (33) with *Siglech*-iCre mice (34). However, even in the absence of Cre expression, BM-derived pDCs from *Batf*-floxed mice exhibited significantly reduced *Batf* expression compared with WT controls (Fig. S10e), indicating that loxP-site insertion dysregulates *Batf* expression in pDCs independently of Cre-mediated recombination. Thus, this mouse line is unsuitable for assessing the specific in vivo contribution of BATF in pDCs. As our analysis was restricted to pDCs, BATF expression in other cell types, including B cells, T cells, and ILCs, may remain unaffected. Subsequently, we analyzed potential direct antiviral effects of BATF-deficient pDCs in an in vitro fibroblast-pDC coculture system. Here, a lower infection rate of fibroblasts was observed in the presence of BATF-deficient pDCs compared to an equivalent number of WT pDCs (Fig. 7c). These findings indicate a modulatory role of BATF in pDCs regarding the IFN-I response, resulting in reduced virus control in the presence of BATF.

In conclusion, our data suggest a model in which low amounts of BATF function to restrain the tonic IFN-I response in resting pDCs, thereby preventing immune pathologies or the development of autoimmunity. Activation of pDCs via PRRs such as TLRs in the case of an infection overrides the BATF-mediated constraint on IFN production for a limited time (Fig. S11a). PRR signaling also ultimately results in increased BATF, leading to downmodulation of pDC activation at later timepoints. Mechanistically, BATF does not dampen IFN-I expression by directly binding to the promoters of IFN-I genes but by binding to the DC-SCRIPT gene and inducing its expression. During the phase of IFN repression, DC-SCRIPT interacts with IRF7, limiting the action of the latter in controlling IFN-I gene expression (Fig. S11b).

Taken together, the results of the present study establish BATF and DC-SCRIPT as two pivotal molecular switches for the regulation of IFN-I responses in pDCs.

## Discussion

The timing and magnitude of IFN-I play a critical role in balancing the promotion of protective immune responses and mitigating immunopathologies (5, 32, 35). Hence, identifying the factors affecting the production of IFN-I and the molecular mechanisms involved are crucial for designing treatments tailored to boost or dampen IFN-I responses. Our data identify BATF as a negative regulator of TLR7/9-induced IFN-I expression in pDCs, acting primarily through DC-SCRIPT rather than direct repression of IFN-I genes, with the exception of *Ifna13* and *Ifna14*.

BATF has previously been associated with the development and function of multiple immune cell lineages (9). However, its expression and function in pDCs have not been characterized. We showed that BATF is expressed at low levels in pDCs; however, following endosomal TLR stimulation, BATF expression is substantially increased. The expression of *Batf* in pDCs trailed behind the expression of *Ifnb1 but* was independent of IFN-I signaling. IFNAR signals, however, may only contribute to *Batf* expression in pDCs in the presence of a second signal induced by TLR9 activation in pDCs. Previous studies have demonstrated the requirement of the TLR4/MyD88-NFκB axis for induction of BATF expression in a B cell line (36–38). In addition, cytokines such as IL-10, LIF, IL-6, IL-7, and IL-21, which signal through Jak/Stat signaling pathways, have been shown to induce BATF in various cell types (39–41). Strikingly, blocking the NFκB activity by specifically inhibiting TAK1 or Stat3 and Stat6 functions abrogated Myd88-mediated BATF expression in large B cell lymphoma cells (38) or IL-7-induced BATF expression in ILC progenitors (14), respectively. In addition, Notch signaling has also been associated with BATF expression in B cells (42). These findings suggest a complex, cell type-specific transcriptional regulation of BATF. Additional investigations are required to identify the transcription factors and the mechanisms involved, including the contributions of other cytokines, such as IL-6 and IL-10, as well as associated signaling pathways, to the expression of BATF in pDCs. Our data define BATF as a negative regulator of TLR7/9-induced IFN-I expression in pDCs. However, BATF neither binds to IFN gene promoter regions, with the exception of *Ifna13* and *Ifna14,* nor directly interferes with IRF7-mediated activation of IFN gene promoters. Instead, BATF indirectly downregulates IFN-I by inducing the expression of DC-SCRIPT in pDCs.

The transcriptional network controlling *Zfp366* expression in pDCs remains unclear. Although PU.1 regulates *Zfp366* in cDCs, the factors cooperating with BATF in pDCs remain to be identified. One could speculate that Spi-B, an ETS family transcription factor highly expressed in pDCs (43, 44) and functionally related to PU.1 (45), may cooperate with BATF to regulate *Zfp366* expression in pDCs. Further studies are needed to define this regulatory network.

Beyond IFN-I, BATF was found to regulate the expression of several cytokines in pDCs. Our data suggest that some cytokines are direct BATF targets, whereas others, including *Il6* and *Il18*, may be regulated indirectly. Further studies are needed to clarify these regulatory mechanisms.

Dysregulated BATF and ZNF366 expression or genetic variants of these genes have been associated with several inflammatory diseases. Both genes show increased transcript levels in peripheral blood cells from septic patients (46, 47), in which pDCs contribute to systemic inflammation (48). GWAS data further link *BATF* and *ZNF366* variants to rheumatoid arthritis (30, 49, 50), a disease in which aberrant IFN-I signaling plays a pivotal role (51, 52). Here, the rheumatoid arthritis-associated variants rs7155603 and rs175714 in *BATF* and rs2278600 in *ZNF366* further support a potential role for the BATF-DC-SCRIPT axis in IFN-I-driven inflammation. While these findings point to a pathophysiological role of *BATF* and *ZNF366* in these inflammatory disorders, experimental proof of the contribution of *BATF* and *ZNF366,* pDCs, and IFN-I to their progression or amelioration remains elusive.

Our data unveiled the interaction of IRF7 with DC-SCRIPT, representing a possible mechanism to inhibit IRF7-induced IFN-I production in pDCs. These data significantly contribute to the understanding of the molecular mechanisms underlying the regulation of IFN-I expression. Mechanistically, DC-SCRIPT may either sequester activated IRF7 or interfere with the binding of IRF7 to the promoter regions of IFN genes. Alternatively, the interaction of IRF7 with DC-SCRIPT might interfere with the recruitment of coactivators or additionally recruit corepressors such as CtBP1 to the transcriptional complex, which is known to physically interact with DC-SCRIPT (22, 53, 54). In addition, the presence of DC-SCRIPT in protein complexes containing the transcription corepressor receptor-interacting protein 140 (RIP140) (53), DNA modifying enzymes such as HDACs (53), and various nuclear receptors (55), including estrogen receptor-α (ER), retinoic acid receptor alpha (RARα) (53, 54), vitamin D receptor (VDR) (56), and glucocorticoid receptor (GR) (57), has been reported, suggesting alternative regulatory mechanisms.

We and others have reported that only a small proportion of pDCs is responsible for the initial expression of IFN-I following systemic activation by TLR7/8/9 stimulation or viral infection (19, 58, 59). During homeostasis, pDCs produce low tonic amounts of IFN-I to keep the transcriptional machinery ready for rapid induction of IFN-I upon pathogen recognition (25, 60). Maintenance of this basal cell state is essential for preventing constitutive autoinflammation in the resting immune system. Exposure to distinct classes of pathogens results in a substantial but transient transcriptomic shift in pDCs, which must be well controlled and reversible to avoid unfavorable immunopathologies (61). Thus, simultaneously with pathogen recognition, anti-inflammatory responses are initiated. Our data demonstrate that BATF acts as a negative feedback regulator of the inflammatory cell state. It contributes significantly to changing the transcriptional profile of pDCs during the transition from a basal to an activated cell state and, subsequently, returning to a steady state condition. A key function of BATF is to promote the expression of DC-SCRIPT, which is, in turn, essential for dampening IFN-I production through the regulation of IRF7 function. These findings introduce BATF and DC-SCRIPT as new target molecules for the development of novel therapeutic interventions aimed at fine-tuning IFN-I levels and controlling IFN-I-induced pathologies.

Although BATF-deficient mice exhibit elevated basal ISG expression, they neither clear viral infections more efficiently in our experimental conditions nor develop overt autoimmune symptoms, potentially reflecting broader effects of BATF deficiency on ILC and T-cell responses (62–65). As BATF and DC-SCRIPT influence multiple hematopoietic lineages, the BATF-DC-SCRIPT axis may also regulate IFN-I responses in other immune or structural cell types, including cDCs, monocytes, macrophages, endothelial cells, epithelial cells, and keratinocytes. Cell type-specific single- and double-knockout models, particularly a pDC-specific *Batf* deficiency, will help define the contributions of this axis to infection and autoimmunity in future studies. Finally, its functional relevance in human pDCs and patient-derived samples must be firmly established before therapeutic strategies targeting BATF or DC-SCRIPT can be considered.

## Materials and Methods

### Animal experiments

*Batf*^-/-^, IFNβ^mob/mob^, *Batf^fl/fl^*, *Siglech*-iCre and *Ifnar*^-/-^ mice have been described previously (12, 33, 34, 66, 67). The mice were maintained in the animal facility of the Heinrich Heine University Düsseldorf, Germany, according to institutional guidelines. *Zfp366*^-/-^ mice and *Zfp366-tdTomato* mice (24) were maintained in the animal facility of the Walter and Eliza Hall Institute, Australia. CD45.1-expressing B6.SJL-*Ptprc^a^ Pepc^b^*/BoyCrl (Ly5.1) mice were purchased from Charles River Laboratories. All mice used in this work were maintained on a C57BL/6 background and housed under SPF conditions. Sex- and age-matched littermates (7-14 weeks old) were used. Animal procedures were approved by the competent authorities of North Rhine-Westphalia, Germany. Where indicated, mice were intravenously infected with 200 PFU of LCMV-WE or injected intravenously with CpG2216 or CpG1668 complexed with DOTAP for the indicated durations. LCMV titers in tissues homogenates were analyzed as described earlier (68).

BM chimeras were generated by transferring 1 x 10^6^ fetal liver cells or 5 x 10^6^ BM cells from CD45.2 *Zfp366*^+/+^ or *Zfp366*^-/-^ donor mice into lethally γ-irradiated (1 x 10.5 Gy) CD45.1 congenic recipient mice. Immune cell populations were examined by flow cytometry 60 days after transfer.

### Cell culture, differentiation and stimulation

293FT, L929 and MC57 cells were cultured in DMEM containing 10% FBS and 2 mM L-glutamine. Hoxb8-FL progenitor lines expressing EGFP or BATF or DC-SCRIPT along with EGFP were generated by lentiviral transduction and maintained in RPMI 1640 supplemented with serum, glutamine, antibiotics, β-mercaptoethanol, β-estradiol and Flt3 ligand (Flt3L). BM-derived pDCs were generated by culturing BM cells in Flt3L-containing medium for seven days. Hoxb8-FL progenitors were differentiated by β-estradiol withdrawal and cultured for seven days. Where indicated, cells were stimulated with CpG2216 (CpG-A), CpG1668 (CpG-B), each complexed with DOTAP, or IFNα4 for the indicated times and concentrations. The sources of cells, detailed culture and stimulation conditions are provided in the relevant figure legends and SI Appendix.

### Flow cytometry and cell sorting

BM cells were obtained from femurs and tibias. Spleens were minced and digested with collagenase and DNase before red-cell lysis. Single-cell suspensions were filtered, Fc receptors were blocked with anti-CD16/CD32, and cells were stained with fluorophore-conjugated antibodies and viability dyes. pDCs were identified as live, single CD3ε^-^, CD19^-^, CD11c^+^, CD11b^-/low^, B220^+^, CD317^+^ and SiglecH^+^ cells, unless stated otherwise. Flow cytometry was performed using FACSCanto II or FACSFortessa instruments and analyzed with FlowJo. Cells were sorted using a FACSAria III. Antibody clones, fluorophores, complete staining panels and procedures are listed in the SI Appendix, Table S8.

### LCMV infection and cytokine measurements

For in vitro infection experiments, MC57 fibroblasts were seeded one day before infection, and FACS-purified WT or *Batf^-/-^* BM-derived pDCs were added 18 h before infection. Cultures were infected with LCMV-WE at a multiplicity of infection of 0.1 for 4 h, and infection was assessed by staining with the VL-4 anti-LCMV antibody.

For cytokine measurements, FACS-purified pDCs were plated and stimulated with CpG2216 or CpG1668 complexed with DOTAP. IFNα, IFNβ and IL-12p40 were measured in culture supernatants or serum by ELISA. A 36-plex ProcartaPlex mouse cytokine/chemokine panel and a Luminex 100 instrument were used for multiplex profiling after 16 h of stimulation. A detailed description of the analysis, stimulation conditions and citations is given in the SI Appendix.

### RT-qPCR

Total RNA was isolated from purified resting or stimulated pDCs or tissue homogenates using the NucleoSpin RNA Mini Kit. cDNA was synthesized using SuperScript III reverse transcriptase and oligo(dT). Quantitative PCR was performed using TaqMan probe-based assays or SYBR Green chemistry, as appropriate, with *Actb* as the reference gene. Detailed methods, primer sequences and probe identifiers are provided in the SI Appendix, Table S9.

### Immunoprecipitation and immunoblotting

For immunoprecipitation, transfected cells were lysed in modified RIPA buffer and incubated with anti-Myc affinity gel overnight. Immunoprecipitates and whole-cell lysates were separated by SDS-PAGE, transferred to nitrocellulose membranes and analyzed by immunoblotting with the indicated antibodies. Method details, antibody and plasmid information are provided in the SI Appendix.

### Plasmids and reporter assay

Expression vectors encoding murine BATF, PU.1 and DC-SCRIPT, as well as IRF7 and TBK1 expression constructs, were used as indicated. Firefly luciferase reporters driven by the *Ifnb1*, *Ifna4* or *Zfp366* promoter were co-transfected into 293FT cells with the indicated expression vectors and a Renilla luciferase normalization plasmid. *Zfp366* promoter deletion constructs containing progressively shorter promoter fragments were used to assess promoter activity. Cells were transfected using polyethyleneimine and incubated for 20 h. Firefly and Renilla luciferase activities were measured using a dual-luciferase assay, and firefly signals were normalized to Renilla activity. Complete construct descriptions and transfection conditions are provided in the SI Appendix.

### RNA-seq, ChIP-seq and ATAC-seq

For RNA-seq, one million FACS-purified BM-derived pDCs were left untreated or stimulated with CpG2216 for 2, 6 or 12 h. RNA quality was assessed before library preparation using the Illumina TruSeq Stranded mRNA Library Prep Kit. Libraries were sequenced on an Illumina HiSeq 3000/4000 platform (single-end, 1 × 150 bp), with three biological replicates per condition. Reads were processed using CLC Genomics Workbench, mapped to the mouse mm10 reference genome, and analyzed for differential expression with edgeR; P values were adjusted for the false discovery rate (FDR).

For BATF ChIP-seq, purified BM-derived pDCs were left untreated or stimulated with CpG2216 for 2 h, crosslinked with formaldehyde and processed for chromatin immunoprecipitation. BATF-associated chromatin was enriched with an anti-BATF antibody, with Drosophila chromatin used for spike-in normalization. Libraries were sequenced on an Illumina NextSeq 500. Reads were aligned to the mm10 and dm3 genomes using BWA; peaks were identified with MACS2, and ENCODE blacklist regions were excluded.

For ATAC-seq, viable purified pDCs from WT and *Batf^-/-^* mice were analyzed at steady state or after 2 h of CpG2216 stimulation. Libraries were prepared by transposase-mediated tagmentation and sequenced on an Illumina NextSeq 500. Reads were aligned to mm10, peaks were called with MACS2, and differential accessibility was analyzed with DESeq2. Methods, processing parameters and quality-control procedures are detailed in the SI Appendix

### Bioinformatics and statistical analysis

Omics data were visualized using R-based plotting tools, Morpheus, eulerr, Cytoscape and Integrative Genomics Viewer, as appropriate. Pathway enrichment used Reactome and gene-set enrichment analysis (GSEA) with hallmark gene sets. Public human expression data (GSE93679) were analyzed using Biobase, GEOquery and limma. Genetic associations at the *BATF* and *ZNF366* loci were obtained from the GWAS Catalog and visualized after conversion of genomic coordinates where required. AnimalTFDB was used for transcription-factor and co-regulator classification. Statistical analyses were performed using GraphPad Prism 9. Student’s t test or one-way or two-way ANOVA with multiple-comparison tests were used as specified in the figure legends. P < 0.05 was considered statistically significant. Analysis details are provided in the SI Appendix

## Supporting information

SI Apendix

## Author Contributions

Conceptualization, S.A., R.M., J.A. and S.S.; software, R.M.; formal analysis, S.A., R.M., P.P., K.K. and S.Z.; investigation, S.A., R.M., M.M., B.L., J.H., R.J.D., M.H.B.,S.Z. and H.C.X.; resources, S.S., S.Z., M.C., S.L.N., J.A. and P.A.L.; writing original draft, S.A., R.M., and S.S.; writing, review & editing, S.A., R.M., R.J.D., K.K.,M.C., S.L.N., P.A.L., J.A., and S.S.; visualization, R.M., S.A., and S.S.; funding acquisition, J.A., S.L.N., M.C., P.A.L. and S.S.; supervision, S.A. and S.S.; all authors reviewed and approved the manuscript.

## Competing Interest Statement

The authors declare no competing interests.

## Acknowledgements

We thank Sonja Schavier, Heinrich Heine University Düsseldorf for her excellent technical support. This work was funded by the German Research Foundation (DFG–SCHE692/6–1) and the Manchot Graduate School ‘Molecules of Infection III and IV’ to SS, the DFG, XU160/3-1 to HCX, the DFG LA2558/8-1, RTG1949 to PAL, the DFG EXC 1003, Grant FF-2014-01 Cells in Motion–Cluster of Excellence, Münster, Germany, and the DFG FOR2107 AL1145/5–2 to JA, the National Health and Medical Research Council of Australia (1155342 and 2000461) to SLN, (1196255) to MC and Cancer Council Victoria, the NHMRC investigator grant (2025614), and The Jack Brockhoff Foundation.to SZ.

## Ethics declarations

All animal procedures in the study were approved by the government of North-Rhine Westphalia, Germany (reference numbers: 81-02.04.2020.A331, 81-02.04.2020.A318, and 84-02.05.30.13.034).

## Resource availability

### Materials availability

The plasmids and cell lines generated in this study are available upon appropriate request.

### Data availability

The generated RNA-seq, ChIP-seq and ATAC-seq data have been deposited in the Gene Expression Omnibus (GEO) and are publicly available as of the date of publication. The accession numbers are listed below.

#### RNA-seq

Next-generation sequencing data of pDC transcriptomes from a longitudinal activation study (CpG 0 h, 2 h, 6 h, 12 h) are available under accession numbers GSE170750 (WT), GSE176419 (*Batf* KO) and GSE176420 (superseries accession number for both WT and KO samples).

ChIP-seq (BATF) data of WT pDCs (CpG 0 h, 2 h) are available under accession number GSE171868.

ATAC-seq data are available under accession numbers GSE171075 (WT pDC; CpG 0 h or 2 h), GSE178410 *(Batf-*KO pDC; CpG 0 h or 2 h), and GSE178412 (superseries accession number for both WT and KO samples).

Published microarray expression-profiling data from human pDCs and murine IFN-β-producing pDCs are available in the GEO under accession number GSE93679 and GSE68788, respectively.

Any additional information required to reanalyze the data reported in this paper is available from the lead contact upon request.

Source data are provided with this paper.

### Code availability

This paper does not report original code.

## Notes

### Competing Interest Statement

The authors have declared no competing interest.

### Summary of Updates

In the revised version of the manuscript, we have updated Figures 1, 6, and 7, along with the corresponding text in the Results section. We have also streamlined the overall manuscript to meet the length requirements and transferred some of the methodological details to the Supporting Information (SI Appendix). The updated submission additionally includes the SI Appendix, comprising Figures S1-S11, their corresponding figure legends, and the Supporting Information Materials and Methods section. Furthermore, we have included seven supplementary data tables as separate Excel files to support the findings presented in the manuscript.

