## Supplementary material for "BATF controls IFN-I production via DC-SCRIPT in plasmacytoid dendritic cells": SI Apendix

[*]Corresponding author: Stefanie Scheu

**This PDF file includes:**

Figures S1 to S11

Legends for Figures S1 to S11

SI: Materials and Methods

Tables S8; S9

SI: References

**Other supporting materials for this manuscript include the following:**

Datasets as supporting information: Tables S1 to S7 in xlxs format as separate files

### Supporting Information: Figures

#### Fig. S1.

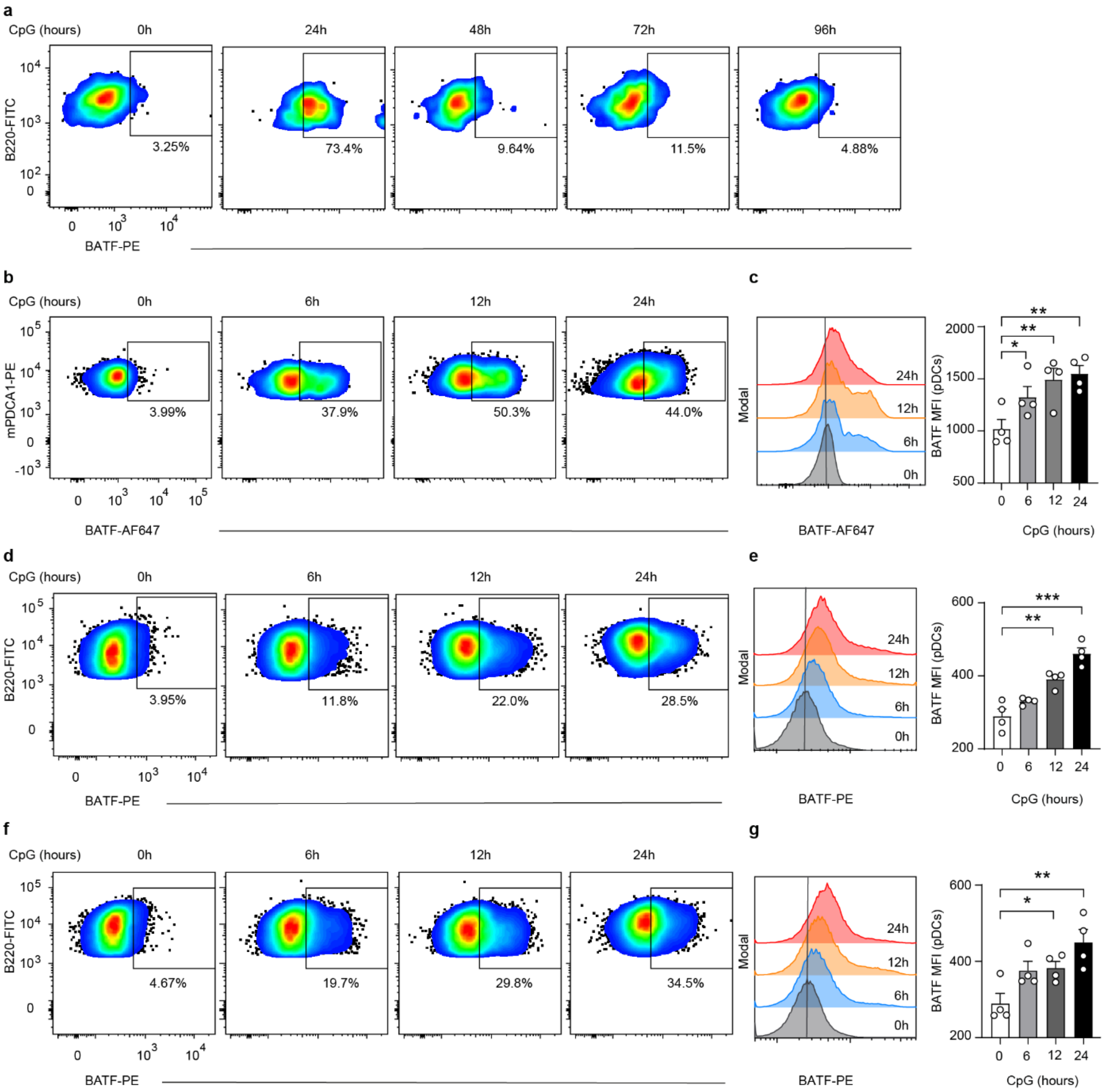


**Fig. S1: TLR9 activation induces BATF expression in pDCs.**
(**a**) Pseudocolor plots showing the expression of BATF in splenic pDCs (CD3$\varepsilon^-$, CD19$^-$, CD317$^{high}$ CD11c$^{int}$, CD11b$^-$ and B220$^+$) from C57BL/6 mice at steady state or after in vivo injection of CpG-A (2216 for the indicated time periods. (**b, c**) Pseudocolor plots (**b**), histograms (**c,** left) and bar chart (**c**, right) showing the expression of BATF in splenic pDCs (CD3$\varepsilon^-$, CD19$^-$, CD317$^{high}$ CD11c$^{int}$ and CCR9$^+$) from C57BL/6 mice at steady state (0 h) or after in vivo injection of CpG-B (1668) for the indicated time durations. (**d-g)** Pseudocolor plots (**d**, **f**), histograms (**e**, **g**, left) and bar chart (**e**, **g**, right) showing expression of BATF in BM-derived Flt3L-cultured pDCs after in vitro stimulation with CpG-A (2216; **d**, **e**) or CpG-B (1668; **f**, **g**) for the indicated durations. The vertical grey line (**c**, **e**, **g**; left) indicates the BATF-median fluorescence intensity (MFI) in resting pDCs. Bar graphs (**c**, **e**, **g**; right) show the MFI (mean $\pm$ s.e.m) of BATF expression in splenic (**c**, right) or BM-derived (**e**, **g**, right) pDCs. Statistical significance was assessed by one-way ANOVA followed by Fisher's LSD multiple comparison test (**c**, **e**, **g**). Data in (**a**) are from a representative sample corresponding to Fig. 1**a**. The data in (**a-g**) represent one representative experiment performed in quadruplicate (n = 4 mice or independent culture per group) out of at least two independent experiments with comparable results. *: $p < 0.05$, **: $p < 0.002$, ***: $p < 0.001$. MFI: median fluorescence intensity; pDCs: plasmacytoid dendritic cells; s.e.m: standard error of the mean.

**Fig. S2.**

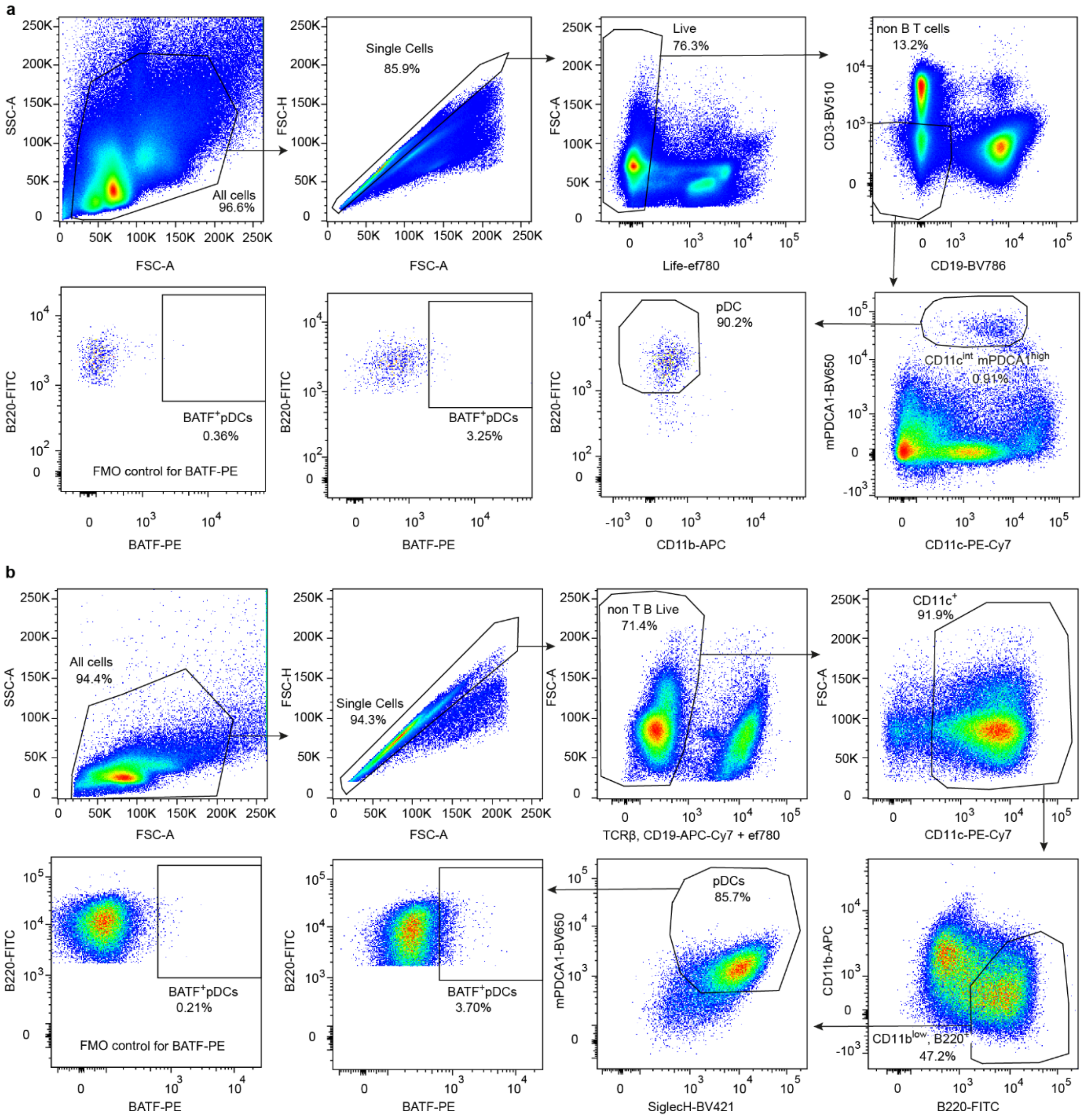


**Fig. S2: Flow cytometry gating strategy for identifying BATF-expressing pDCs.**
(**a**) Splenocytes from C57BL/6 mice were stained and first gated based on FSC-A and SSC-A. After exclusion of doublets, live cells (ef780$^-$) were selected and further gated to exclude lymphocyte lineages (CD3ε$^-$ CD19$^-$). pDCs were identified as CD11b$^-$ B220$^+$ cells pre-gated for high mPDCA-1 (CD317) and intermediate CD11c expression. (**b**) Flow cytometry gating strategy for identifying BM-derived pDCs in Flt3L cultures after stimulation with CpG. Cell-sized particles were gated based on FSC-A and SSC-A, followed by exclusion of doublets. Live (ef780$^-$) non-B (CD19$^-$) and non-T (TCRβ$^-$) cells were analyzed for CD11c expression. CD11c$^+$ cells were further gated for low to negative CD11b expression and positive B220 expression. pDCs were identified as CD11b$^{low}$ B220$^+$ cells that were additionally positive for mPDCA-1 (CD317) and Siglec-H.

**Fig. S3**

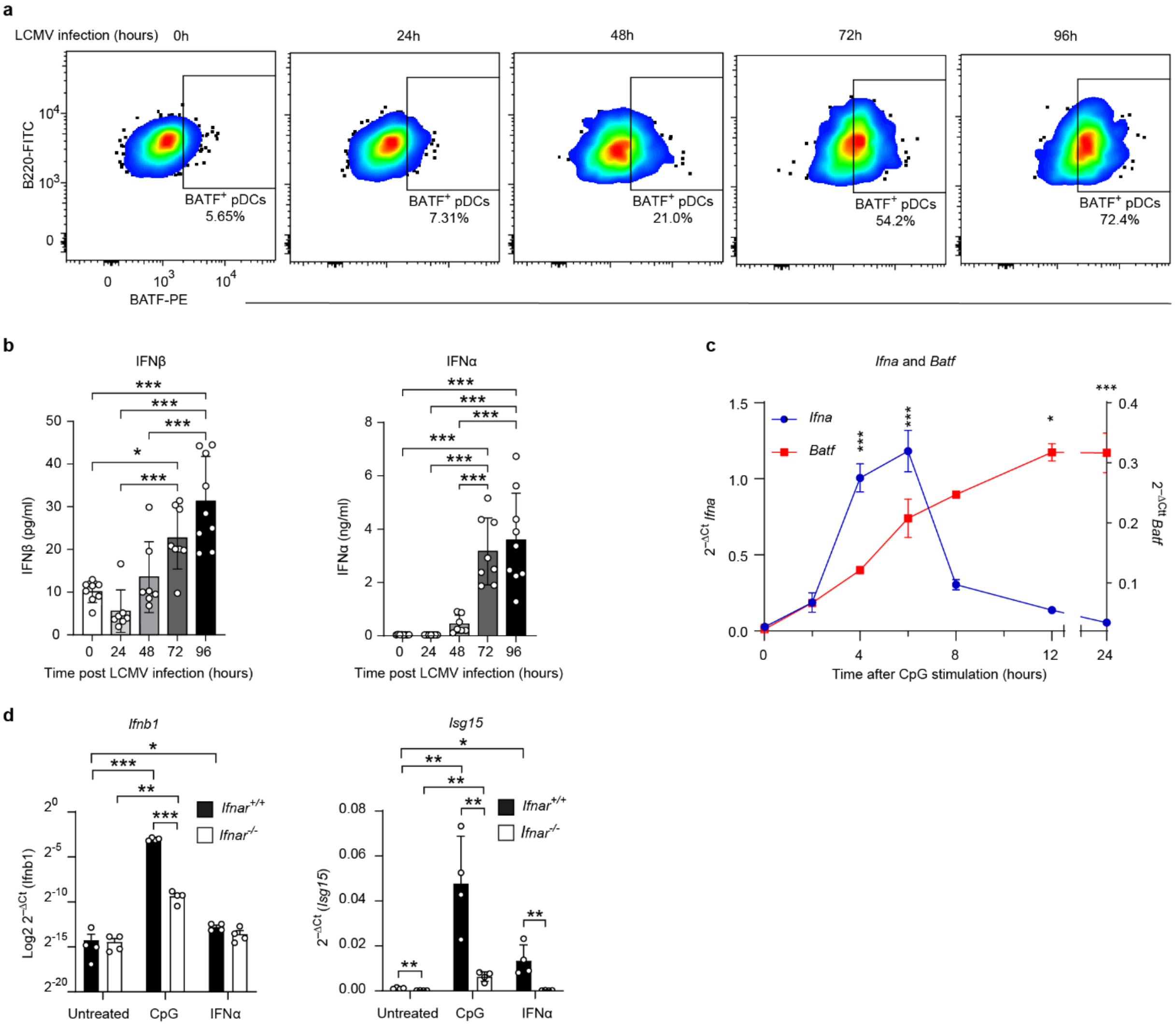


**Fig. S3: LCMV infection induces BATF expression in pDCs.**

(a) Pseudocolor plots showing BATF expression in splenic pDCs ($CD3\varepsilon^-$ $CD19^-$ $CD317^{high}$ $CD11c^{int}$ $CD11b^-$ $B220^+$) from C57BL/6 mice at steady state (0 h) or after systemic LCMV infection for the indicated time periods. (**b**) Bar graphs showing serum levels of IFNβ (left) and IFNα (right) after systemic LCMV infection for the indicated time periods. (**c**) Quantitative qRT-PCR analysis showing relative expression of *Ifna* (multiple *Ifna* genes amplified by wobble primers, blue line) and *Batf* (red line) mRNA in FACS-purified murine BM-derived Flt3L-cultured pDCs analyzed at resting state (0 h) or after activation with CpG for indicated durations. (**d**) Quantitative RT-PCR analysis showing relative expression of *Ifnb1* (left) and *Isg15* (right) mRNAs in BM-derived Flt3L-cultured, FACS-purified pDCs from *Ifnar*$^{-/-}$ and wild-type (*Ifnar*$^{+/+}$) mice at steady state (untreated) or after 6 h stimulation with 1 μM CpG or 100 units/ml IFNα4. Data represent means ± s.e.m. and were normalized to *Actb*. Data in **a** are from representative sample corresponding to Fig. 1**b**. Data in (**b**) are pooled data from two independent experiments. Data in (**c**) are from one representative experiment out of three independent experiments with comparable results. Significant differences were analyzed by one-way ANOVA followed by multiple t test with Bonferroni correction (**c**), or by two-way ANOVA followed by multiple comparisons tests with Bonferroni correction (**d**). Gates in (**a**) were applied using FMO controls. *: $p < 0.05$, **: $p < 0.002$, ***: $p < 0.001$. pDC: plasmacytoid dendritic cell; s.e.m.: standard error of the mean; FMO: fluorescence minus one.

**Fig. S4**

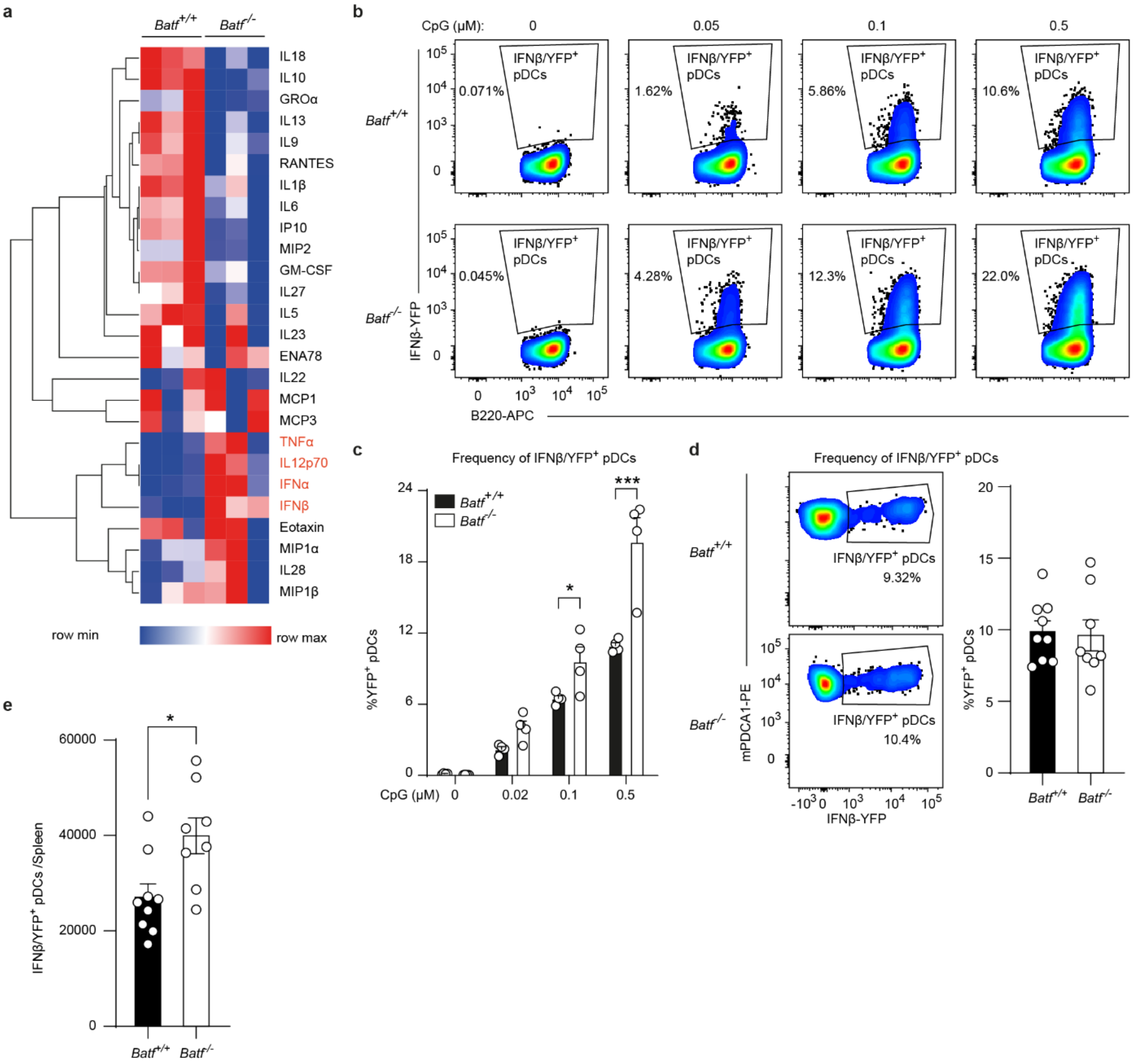


### Fig. S4: BATF influences the expression of various cytokines in pDCs.

Heatmap showing the cytokine/chemokine expression profile of CpG (1 μM for 16 h)-stimulated, FACS-sorted BM-derived Flt3L-cultured pDCs from *Batf*$^{-/-}$ and WT (*Batf*$^{+/+}$) mice. The quantities of measurable analytes quantified by a multiplex cytokine array are shown in the heatmap (n = 3 per group). Data shown are from one experiment performed in triplicate. (**b**, **c**) Pseudocolor plots, (**b**) and bar chart (**c**) showing IFNβ/YFP-expressing BM-derived Flt3L-cultured pDCs from wild type and *Batf*-deficient IFNβ/YFP reporter (MOB) mice following in vitro stimulation with the indicated concentrations of CpG-A (2216) for 4 hours. Shown are frequency of IFNβ/YFP$^{+}$ pDCs. Data in **b** represent a sample shown in the quantitative summary in (**c**). (**d**, **e**) Pseudocolor plots, (**d**, left) and bar chart (**d**, right, **e**) showing IFNβ/YFP-expressing pDCs analyzed ex vivo from wild type (*Batf*$^{+/+}$) and *Batf*-deficient (*Batf*$^{-/-}$) IFNβ/YFP reporter (MOB) mice 6 h after in vivo injection of CpG-A (2216). Shown are the frequency of IFNβ/YFP$^{+}$ pDCs (**d**), and the absolute number of IFNβ/YFP$^{+}$ pDCs (**e**). Pseudocolor plots in (**d**, left) represent samples included in the quantitative summary shown in (**d**, right). Data in (**b**, **c**) represent one experiment performed in quadruplicate (4 independent cultures per group) and are representative of at least two independent experiments with comparable results. Data in (**d**, **e)** represent pooled results from 8-9 mice per group across two independent experiments. Statistical significance was determined using two-way ANOVA followed by Fisher's LSD multiple-comparison test (**c**) or unpaired two-tailed *t*-test with Welch's correction (**d**, **e**). *: $p < 0.05$, **: $p < 0.002$, ***: $p < 0.001$. MFI: median fluorescence intensity; pDCs: plasmacytoid dendritic cells; s.e.m.: standard error of the mean; MOB: messenger of IFNβ.

**Fig. S5**

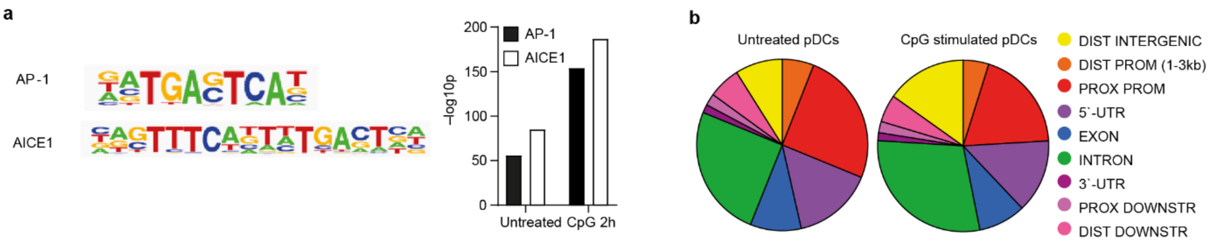


**c**

Homer de novo Motif Results (Batf $^{+/+}$ Untreated)

| Rank | Motif | P-value | % of Targets | Best Match/Details |
|---|---|---|---|---|
| 1 | TCATAATGAAAC | 1e-91 | 10.26% | IRF:BATF(IRF:bZIP)/pDC-Irf8-ChIP-Seq(GSE66899)/Homer(0.911) |
| 2 | TATGAGTCAT | 1e-68 | 12.01% | BATF(bZIP)/Th17-BATF-ChIP-Seq(SE39756)/Homer(0.971) |
| 3 | AAAGAGGAAGTG | 1e-33 | 9.50% | SpiB(ETS)/OCILY3-SPIB-ChIP-Seq(GSE56857)/Homer(0.956) |
| 4 | GTTATGACTG | 1e-19 | 2.51% | MAFG::NFE2L1/MA0089.1 /Jaspar(0.700) |
| 5 | TGTGGTTAAC | 1e-16 | 9.93% | RUNX-AML(Runt)/CD4+-PolII-ChIP-Seq(Barski_et_al.)/Homer(0.846) |
| 6 | CAGAAACCGC | 1e-15 | 19.76% | RUNX2/MA0511.2/Jaspar(0.726) |
| 7 | GTGACGTCACCC | 1e-14 | 6.44% | Atf7(bZIP)/3T3L1-Atf7-ChIP-Seq(GSE56872)/Homer(0.939) |
| 8 | GCGGAGGGATAC | 1e-13 | 1.31% | ZNF467(Zf)/HEK293-ZNF467.GFP-ChIP-Seq(GSE58341)/Homer(0.606) |
| 9 | GGCCTCGAGA | 1e-13 | 9.28% | ZNF711(Zf)/SHSY5Y-ZNF711-ChIP-Seq(GSE20673)/Homer(0.734) |
| 10 | ATTACGGACG | 1e-13 | 7.21% | HINFP/MA0131.2/Jaspar(0.659) |

**d**

Homer Known Motif Enrichment Results (Batf $^{+/+}$ Untreated)

| Rank | Motif | Name | P-value |
|---|---|---|---|
| 1 | CTTTCATTATGACTC | IRF:BATF(IRF:bZIP)/pDCIrf8-ChIP-Seq(GSE66899)/Homer | 1e-109 |
| 2 | CAGTTTCATTTTGACTCA | bZIP:IRF(bZIP,IRF)/Th17-BatF-ChIP-Seq(GSE39756)/Homer | 1e-85 |
| 3 | GATGAGTCAT | BATF(bZIP)/Th17-BATFChIP-Seq(GSE39756)/Homer | 1e-56 |
| 4 | GATGACTCATCC | Atf3(bZIP)/GBM-ATF3-ChIP-Seq(GSE33912)/Homer | 1e-55 |
| 5 | GAATGAGTCATC | Fra1(bZIP)/BT549-Fra1-ChIP-Seq(GSE46166)/Homer | 1e-52 |
| 6 | GATGAGTCATCC | Fosl2(bZIP)/3T3L1-Fosl2-ChIP-Seq(GSE56872)/Home | 1e-47 |
| 7 | ATGACTCATC | AP-1(bZIP)/ThioMac-PU.1-ChIP-Seq(GSE21512)/Homer | 1e-47 |
| 8 | GATGACTCATCC | Jun-AP1(bZIP)/K562-cJun-ChIP-Seq(GSE31477)/Homer | 1e-45 |
| 9 | TGCTGAGTCA | Bach2(bZIP)/OCILy7-Bach2-ChIP-Seq(GSE44420)/Homer | 1e-22 |
| 10 | AAAGAGGAAGTG | SpiB(ETS)/OCILY3-SPIBChIP-Seq(GSE56857)/Homer | 1e-20 |

**e**

Homer de novo Motif Results (Batf $^{+/+}$ CpG 2h)

| Rank | Motif | P-value | % of Targets | Best Match/Details |
|---|---|---|---|---|
| 1 | TATGAGTCAT | 1e-155 | 27.84% | BATF(bZIP)/Th17-BATF-ChIP-Seq(GSE39756)/Homer(0.986) |
| 2 | GTCATAATGAAA | 1e-128 | 17.26% | IRF:BATF(IRF:bZIP)/pDC-Irf8-ChIP-Seq(GSE66899)/Homer(0.913) |
| 3 | AAGCGGAAGT | 1e-70 | 25.28% | PB0058.1_Sfpi1_1/Jaspar(0.894) |
| 4 | AACCACAA | 1e-36 | 21.71% | RUNX1(Runt)/Jurkat-RUNX1-ChIP-Seq(GSE29180)/Homer(0.958) |
| 5 | GTCACGTCACCG | 1e-26 | 12.81% | CRE(bZIP)/Promoter/Homer(0.876) |
| 6 | CGAAATGA | 1e-20 | 9.69% | PH0037.1_Hdx/Jaspar(0.696) |
| 7 | CGCAGATG | 1e-18 | 37.19% | Ascl1(bHLH)/NeuralTubes-Ascl1-ChIP-Seq(GSE55840)/Homer(0.912) |
| 8 | GGAAATTCCC | 1e-17 | 10.80% | NFkB-p65-Rel(RHD)/ThioMac-LPS-Expression(GSE23622)/Homer(0.993) |
| 9 | CAGTCATAGCTG | 1e-15 | 3.56% | MAFG::NFE2L1/MA0089.1 /Jaspar(0.637) |
| 10 | TCCTCGCCAC | 1e-15 | 4.23% | PB0164.1_Smad3_2/Jaspar(0.696) |

**f**

Homer Known Motif Enrichment Results (Batf $^{+/+}$ CpG 2h)

| Rank | Motif | Name | P-value |
|---|---|---|---|
| 1 | GATGAGTCAT | BATF(bZIP)/Th17-BATF-ChIP-Seq(GSE39756)/Homer | 1e-150 |
| 2 | GATGACTCATCC | Atf3(bZIP)/GBM-ATF3-ChIP-Seq(GSE33912)/Homer | 1e-146 |
| 3 | GAATGAGTCATC | Fra1(bZIP)/BT549-Fra1-ChIP-Seq(GSE46166)/Homer | 1e-143 |
| 4 | ATGACTCATC | AP-1(bZIP)/ThioMac-PU.1-ChIP-Seq(GSE21512)/Homer | 1e-130 |
| 5 | CTTTCATTATGACTC | IRF:BATF(IRF:bZIP)/pDC-Irf8-ChIP-Seq(GSE66899)/Homer | 1e-130 |
| 6 | GATGAGTCATCC | Fosl2(bZIP)/3T3L1-Fosl2-ChIP-Seq(GSE56872)/Homer | 1e-125 |
| 7 | CAGTTTCATTTTGACTCA | bZIP:IRF(bZIP,IRF)/Th17-BatF-ChIP-Seq(GSE39756)/Homer | 1e-119 |
| 8 | GATGACTCATCC | Jun-AP1(bZIP)/K562-cJun-ChIP-Seq(GSE31477)/Homer | 1e-104 |
| 9 | TGCTGAGTCA | Bach2(bZIP)/OCILy7-Bach2-ChIP-Seq(GSE44420)/Homer | 1e-59 |
| 10 | AAAGAGGAAGTG | SpiB(ETS)/OCILY3-SPIB-ChIP-Seq(GSE56857)/Homer | 1e-44 |

**Fig. S5: BATF binds predominantly to proximal promoter and intron regions.**
(**a**) HOMER known motif analysis (JASPAR) for gene regions in untreated and 2 h CpG-A-stimulated, FACS-sorted BM-derived Flt3L-cultured pDCs from C57BL/6 mice that interact with BATF according to ChIP-seq. (**b**) Genomic location distribution of BATF binding sites in untreated or CpG-stimulated pDCs according to BATF ChIP-seq. Two biological replicates were analyzed per condition, and data from pooled samples are shown.(**c**-**f**) Top 10 Homer de novo (**c**, **e**) and known (**d**, **f**) motif enrichment results from anti-BATF ChIP-seq performed on FACS-purified pDCs at steady state (untreated; **c**, **d**) or after activation (2 h CpG stimulation; **e, f**). Data shown in (**c**-**f**) represent one representative sample from two independent biological replicates per condition.

**Fig. S6**

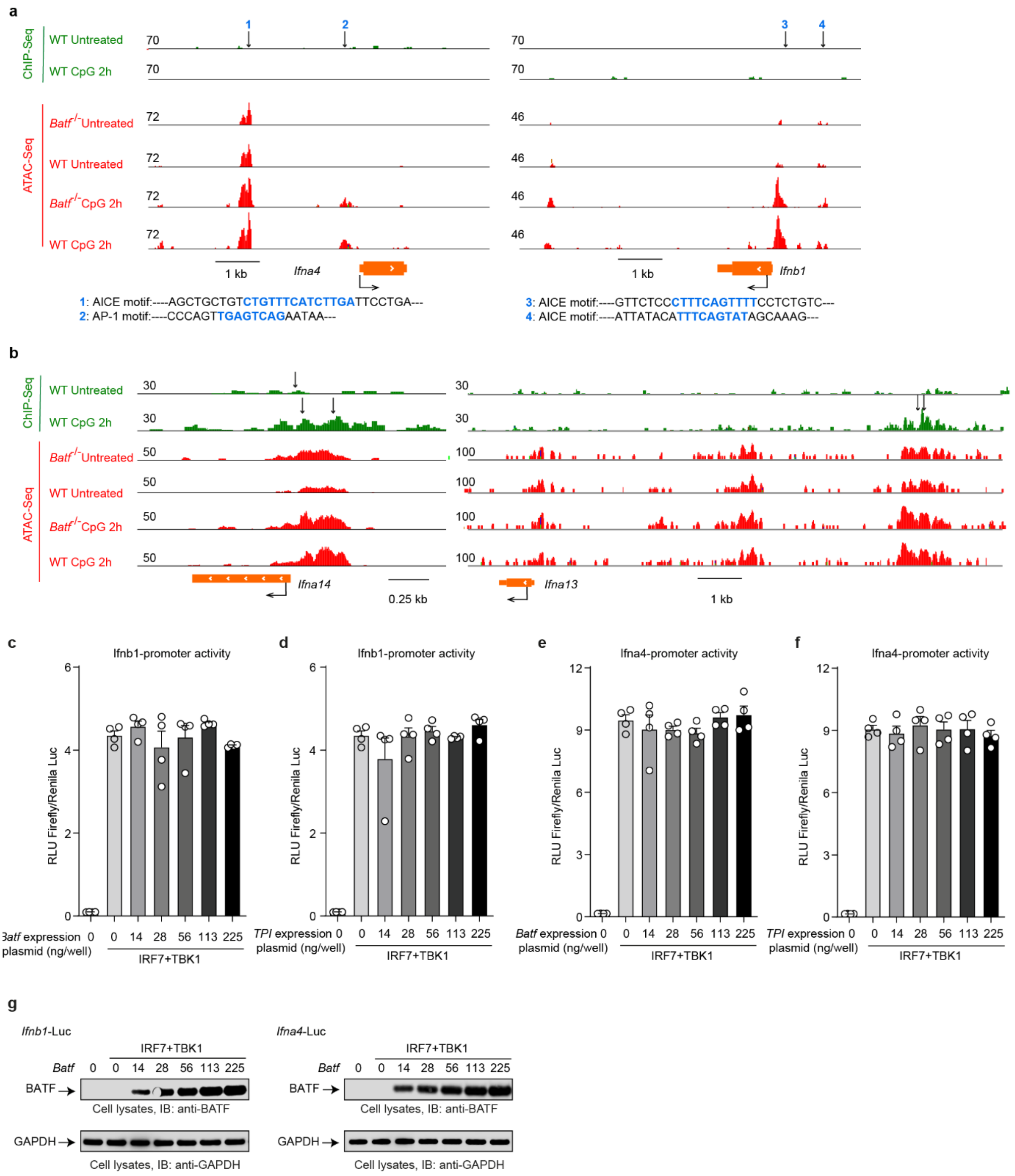


**Fig. S6: BATF binds selectively to the *Ifna13* and *Ifna14* promoter regions amongst all IFN I genes.**

(**a, b**) BATF ChIP-seq (green) and ATAC-seq (red) peaks visualized for the *Ifna4* (left) and *Ifnb1* (right) genes (**a**) and the *Ifna13* (right) and *Ifna14* (left) genes (**b**) using the IGV program. Locations of AICE and AP-1 motifs are indicated by arrows (**a**) and numbered (1-4) within *Ifna4* and *Ifnb* promoter regions. The corresponding AICE and AP-1 sequence (highlighted in blue) is shown below. The track height is shown by the number on the left side. Arrows (**b**) indicate significant BATF binding sites as calculated by the MACS algorithm. (**c-f**) Dual reporter gene assays showing the influence of BATF overexpression on IRF7-induced *Ifnb1* (**c**, **d**) and *Ifna4* (**e**, **f**) promoter activity. *Ifnb1* and *Ifna4* reporter gene assays were performed in 293FT cells. To mimic pDC-like IRF7 activation, IRF7 and TBK1 were overexpressed, which activated *Ifnb1* and *Ifna4* promoters. The housekeeping gene *TPI1* served as a control (**d**, **f**). The amounts of BATF encoding plasmid (ng), used for transfection are indicated below the corresponding bars. (**g**) Western blot images showing BATF expression (upper panel) and GAPDH, as loading control (lower panel). Western blots were perform using pooled cell lysates from the biological replicates used in the dual reporter gene assays shown in (**c**, **e**) for the *Ifnb1*(left) and *Ifna4* (right) promoters. Data shown are from one representative experiment performed in quadruplicate out of three independent experiments with comparable results. Data shown in (**c**-**f**) are the means of the ratios of light units of firefly luciferase/Renilla luciferase ± s.e.m. Significance was analyzed by one-way ANOVA followed by Bonferroni's multiple comparison test. s.e.m.: standard error of the mean.

**Fig. S7**

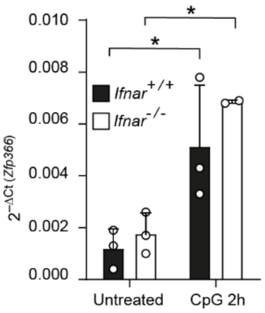


**Fig. S7: IFN-I responses are dispensable for *Zfp366* expression in pDCs.**
Quantitative RT-PCR analysis of *Zfp366* expression in FACS-sorted, untreated or 2 h CpG-A-stimulated WT and *Ifnar*$^{-/-}$ BM-derived Flt3L-cultured pDCs. Data are presented as means ± s.e.m. of relative expression from three biological replicates (normalized to *Actb*). Statistical significance was determined using two-way ANOVA followed by Bonferroni's multiple comparison test. *: $p < 0.05$. s.e.m.: standard error of the mean.

**Fig. S8**

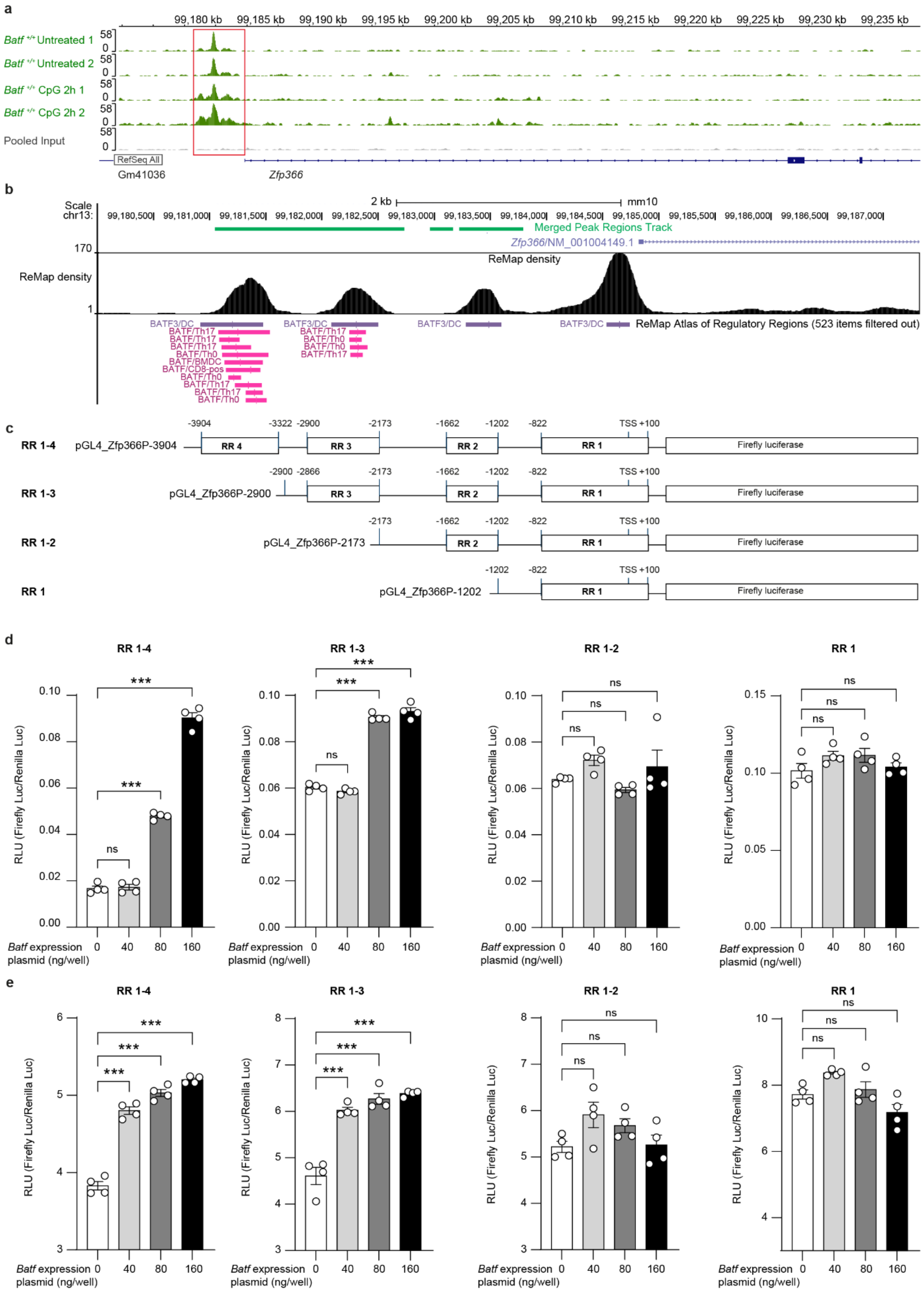
a
Batf +/+ Untreated 1
Batf +/+ Untreated 2
Batf +/+ CpG 2h 1
Batf +/+ CpG 2h 2
Pooled Input
RefSeq All
Gm41036
Zfp366
b
Scale
chr13:
2 kb
mm10
Merged Peak Regions Track
Zfp366/NM_001004149.1
ReMap density
ReMap Atlas of Regulatory Regions (523 items filtered out)
c
RR 1-4 pGL4_Zfp366P-3904
RR 1-3 pGL4_Zfp366P-2900
RR 1-2 pGL4_Zfp366P-2173
RR 1 pGL4_Zfp366P-1202
Firefly luciferase
d
RR 1-4
RR 1-3
RR 1-2
RR 1
RLU (Firefly Luc/Renilla Luc)
Batf expression plasmid (ng/well)
e
RR 1-4
RR 1-3
RR 1-2
RR 1
RLU (Firefly Luc/Renilla Luc)
Batf expression plasmid (ng/well)

**Fig. S8: BATF binds to the promoter of *Zfp366* and promotes its activity.**
(**a**) Integrative Genomics Viewer (IGV) screenshot of anti-BATF ChIP-seq data showing differential binding peaks for binding of BATF to *Zfp366* promoter in all replicates. The track height was adjusted using the "group autoscale" option (numbers on the left). Upper two green tracks: steady state (Untreated 1 and 2); lower two green tracks: activated condition (CpG 2 h 1 and 2); lowest grey track: background signal from pooled input control. **(b)** UCSC genome browser screenshot (mouse GRCm38/mm10) highlighting the *Zfp366* promoter region. Overlapping BATF binding regions from anti-BATF ChIP-seq in pDCs (green tracks, merged peak regions) and previously published data sets from the ReMap atlas for BATF family members are shown. Black peaks (ReMap density) represent four regulatory regions in the *Zfp366* promoter.**(c)** Schematic representation of *Zfp366* reporter gene plasmids (pGL4 vector backbone) showing four regulatory regions (RR1-4) in the *Zfp366* promoter and corresponding deletion mutants. Numbers indicate the positions of regulatory regions with reference to the transcription start site (TSS; upstream "-", downstream "+"). (**d**, **e**) Bar charts presenting the results of dual reporter gene assays showing the influence of BATF overexpression on *Zfp366*-promoter activity. *Zfp366* reporter gene assays were performed in 293FT cells using full-length (RR1-4) or deletion mutants for regulatory regions (RR1, RR1-2, and RR1-3) reporter vectors (shown in **c**) with (**d**) or without (**e**) PU.1 (Spi1) overexpression. The quantities of BATF encoding plasmid (ng), used for transfection, are indicated below the respective bars. Data shown in **d** and **e** are from one representative experiment performed in quadruplicate out of two (**d**) or three (**e**) independent experiments with comparable results. Data represent one experiment performed in quadruplicate, representative of two (**d**) or three (**e**) independent experiments with comparable results. Values are means ± s.e.m. of firefly/Renilla luciferase ratios. Statistical significance was determined by one-way ANOVA with Bonferroni's multiple-comparison test. ***: $p < 0.001$. DC: dendritic cells; s.e.m.: standard error of the mean.

**Fig. S9**

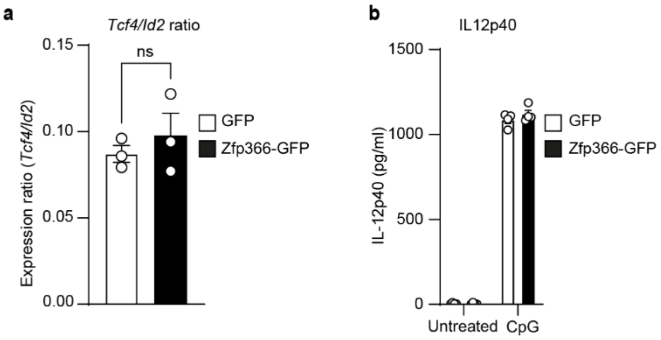


**Fig. S9: DC-SCRIPT overexpression does not influence IL-12p40 expression in pDCs.**
(**a**) Bar charts showing the ratio of *Tcf4* (E2-2) to *Id2* transcripts analyzed by quantitative-RT-PCR in FACS purified Hoxb8-Fl-pDCs overexpressing GFP or ZFP366 and GFP. (**b**) Bar chart presenting IL-12p40 levels in the supernatants of Hoxb8-Fl-pDCs overexpressing GFP or ZFP366 and GFP at steady state (untreated) or following overnight stimulation with CpG-A (2216). Data shown are means ± s.e.m. Data presented are ratios of means (**a**) of relative expression from three biological replicates (normalized to *Actb*). Data shown in (**b**) are the quantities of IL-12p40 analyzed by ELISA and are representative of three independent experiments. Statical differences were tested by students t test (**a**) or two-way ANOVA followed by Bonferroni's multiple comparison test (**b**).

**Fig. S10**

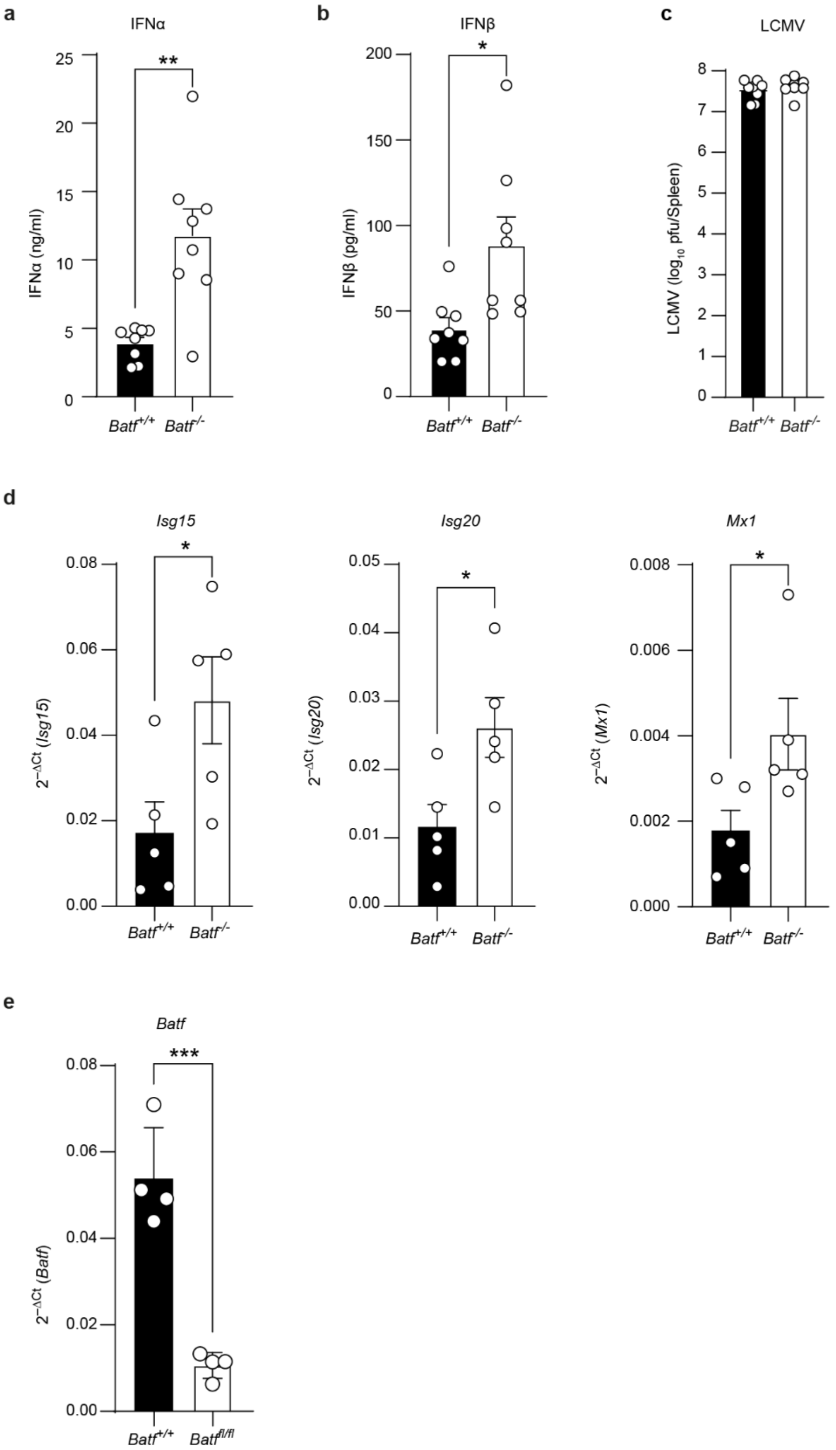


**Fig. S10: BATF-deficient mice exhibit heightened IFN-I and ISGs expression.**

(**a**, **b**) Bar charts showing the serum levels of IFNα (**a**) and IFNβ (**b**) in wild-type (*Batf*$^{+/+}$) and *Batf*-deficient (*Batf*$^{-/-}$) mice 96 h after LCMV infection (200 PFU per mouse), measured by ELISA. (**c**) LCMV titers in the spleen of *Batf*$^{+/+}$ and *Batf*$^{-/-}$ mice, determined by PFU assay. (**d**) Quantitative-RT-PCR analysis of ISGs transcripts (*Isg15*, left bars; *Isg20*, middle bars, and *Mx1,* right bars) in pooled peripheral lymph nodes (inguinal, brachial, and axillary from untreated mice (**e**) Quantitative RT-PCR showing *Batf* expression in FACS purified, BM-derived pDCs from wild-type (*Batf*$^{+/+}$, black bars) and *Batf*-floxed (*Batf*$^{fl/fl}$, white bars) mice. Data in (**d**) represent pooled results from 5 mice, while (**a**-**c**) represent pooled data from 8 mice collected across two independent experiments. Data in (**e**) represent four independent biological replicates (four mice per genotype). Data in (**a**-**c**) were obtained from the same set of mice. Statistical significance was determined by unpaired two-tailed t test (**a**-**e**). *: $p < 0.05$, **: $p < 0.002$. ***: $p < 0.001$

**Fig. S11**

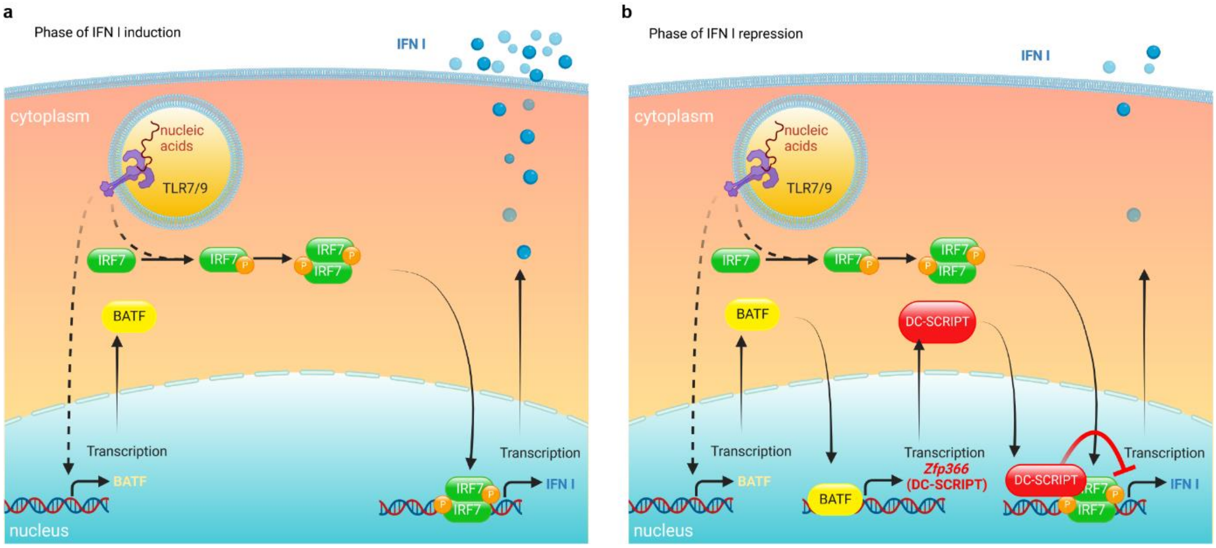


**Fig. S11: BATF dampens IFN I expression via regulation of ZFP366 expression in pDCs.**
(**a, b**) Schematic diagram illustrating the molecular mechanism underlying the BATF-mediated repression of IFN I in pDCs. (**a**) Nucleotide-induced endosomal TLR activation initiates IRF7 phosphorylation, dimerization and nuclear translocation. During the phase of IFN induction, phosphorylated IRF7 binds to consensus target sequences in the promoter regions of IFN I genes and induces massive amounts of IFN I secretion. At the beginning of the phase of IFN induction, minute amounts of BATF are present in the cells, which is not sufficient to repress the strong IRF7 signal. (**b**) In addition to IRF7 activation, endosomal TLR activation also generates a second signal, which leads to the enhanced expression of BATF in pDCs. Elevated BATF protein translocates to the nucleus and induces the expression of DC-SCRIPT by directly binding to the promoter of the DC-SCRIPT gene. During the phase of IFN I repression, newly produced DC-SCRIPT in higher amounts physically interacts with IRF7 and dampens IRF7-induced IFN I production. The diagram was created with BioRender.com.

**Dataset S1 (separate file).**

Table S1. Interferome database genes identified in the differential expression analysis of wild-type (WT) and *Batf*-deficient (KO) pDCs; related to Fig. 3d.

**Dataset S2 (separate file).**

Table S2. Expression of TLR and IFN-I signaling pathway genes in WT and *Batf*-deficient (KO) pDCs (related to Fig. 3d).

**Dataset S3 (separate file).**

Table S3. Expression of IRFs, IFN-I and selected ISGs in the differential expression analysis of wild-type (WT) and *Batf*-deficient pDCs (KO) and BATF ChIP-seq data; related to Fig. 4a.

**Dataset S4 (separate file).**

Table S4. $Log_2$-normalized counts per million (CPM) of differentially expressed transcription factors and co-regulators in WT and *Batf*-deficient (KO) pDCs.

**Dataset S5 (separate file).**

Table S5. RPKM counts of all genes in wild-type (WT) and *Batf*-deficient (KO) pDC samples at steady state and after CpG stimulation (2, 6, 12h).

**Dataset S6 (separate file).**

Table S6. Log2 fold changes and FDR values for all genes in wild-type (WT) and *Batf*-deficient (KO) pDC samples at steady state and after CpG stimulation (2, 6, 12h).

**Dataset S7 (separate file).**

Table S7. *BATF*- and *ZNF366*-associated GWAS results from the NHGRI-EBI GWAS Catalog (https://www.ebi.ac.uk/gwas/); Related to Fig. 6c, d.

### Supporting information: Materials and Methods

#### Mouse strains, animal procedures and bone-marrow chimeras

*Batf*$^{-/-}$, IFNβ$^{mob/mob}$, *Batf*$^{fl/fl}$, *Siglech*-iCre and *Ifnar*$^{-/-}$ mice have been described previously (1-5). The mice were maintained in the animal facility of the Heinrich Heine University Düsseldorf, Germany, according to institutional guidelines. *Zfp366*$^{-/-}$ and *Zfp366-tdTomato* (6) mice were maintained in the animal facility of the Walter and Eliza Hall Institute, Australia. CD45.1-expressing B6.SJL-Ptprc$^{a}$ Pepc$^{b}$/BoyCrl (Ly5.1) congenic mice were obtained from Charles River Laboratories. Mice were maintained on a C57BL/6 background under specific pathogen-free conditions; sex- and age-matched littermates aged 7-14 weeks were used. Animal procedures were approved by the competent authorities of North Rhine-Westphalia, Germany.

For BM chimeras, $1 \times 10^6$ fetal liver cells or $5 \times 10^6$ BM cells from CD45.2 *Zfp366*$^{+/+}$ or *Zfp366*$^{-/-}$ donors were transferred intravenously into lethally γ-irradiated (10.5 Gy) CD45.1 congenic recipients. Recipients were maintained under specific pathogen-free conditions, and immune-cell populations were analyzed by flow cytometry after 60 days.

Where indicated, mice were infected intravenously with 200 PFU of LCMV strain WE for the indicated time spans. For analysis of serum IFN-I levels and ex vivo BATF expression, mice were injected intravenously with 2 nmol CpG2216 or CpG1668 complexed with 30 µg DOTAP in HBSS and analyzed after 6 h or as specified in the relevant experiment.

#### Cell lines and lentiviral transduction

293FT cells were obtained from Invitrogen. L929 and MC57 cells are commercially available from ATCC. These cell lines were cultured in DMEM (Gibco/Thermo Scientific) supplemented with 10% FBS and 2 mM L-glutamine (Sigma-Aldrich/Merck) at 37°C in a humidified incubator with 10% CO2 and passaged twice weekly.

Hoxb8-FL (7) progenitors were provided by Dr. Susanne Kirschnek (University Hospital Freiburg, Germany). Hoxb8-FL-GFP cells expressed EGFP alone, whereas Hoxb8-FL-Batf-GFP and Hoxb8-FL-Zfp366-GFP cells expressed murine BATF or DC-SCRIPT, respectively, together with EGFP. Cells were maintained in low-endotoxin RPMI 1640 (PAN-Biotech) containing 10% FBS (Sigma-Aldrich/Merck), 2 mM L-glutamine, 100 U/ml penicillin, 100 µg/ml streptomycin, 50 µM β-mercaptoethanol (all from Gibco/Thermo Scientific), 1 µM β-estradiol (Sigma-Aldrich/Merck), and either 70 ng/ml recombinant Flt3L (R & D Systems) or 5% conditioned medium from a Flt3L-producing CHO cell line (reported to be approximately equivalent to 70 ng/ml recombinant Flt3L). Cultures were maintained at $1 \times 10^5$-$1 \times 10^6$ cells/ml.

For lentivirus production, 293FT cells were transiently transfected with the pLP/VSVG and psPAX2 plasmids, encoding viral envelope proteins, along with pWPI-based bicistronic lentiviral expression vectors encoding EGFP (pWPI), BATF and EGFP (pWPI-mBatf-GFP), or DC-SCRIPT and EGFP (pWPI-mZfp366-GFP) using JetPRIME transfection reagent (Polypus). Viral supernatants were filtered through a 0.45-µm membrane. Target cells were transduced by centrifugation at 500 × g for 90 min at 30°C in the presence of 4 µg/ml polybrene. The following day, cells were washed and replated in fresh medium. After at least three passages, live GFP-positive cells were purified by FACS using a FACSAria III (BD Biosciences) and expanded.

**Generation of BM-derived dendritic cells and cell stimulation**

BM cells were isolated from femurs and tibias as described earlier (8), subjected to red-cell lysis for 3 min at room temperature, and filtered through a 100-µm cell strainer. Cells were cultured in low-endotoxin RPMI 1640 (PAN-Biotech) containing 10% FBS (Sigma-Aldrich/Merck), 2 mM L-glutamine, 100 U/ml penicillin, 100 µg/ml streptomycin, and 50 µM β-mercaptoethanol (Gibco/Thermo Scientific), supplemented with 5% conditioned medium from a Flt3L-producing CHO cell line (reported to be approximately equivalent to 70 ng/ml recombinant Flt3L). BM-derived Flt3L-cultured DCs were generated as previously described (3). Briefly, cells were plated at $2 \times 10^6$/ml and differentiated for 7 days. On day 5, half of the medium was replaced with fresh differentiation medium containing Flt3L.

Hoxb8-FL progenitor-derived cells were expanded in culture and differentiated as previously described (7). Briefly, Hoxb8-FL progenitors were washed three times with PBS to remove β-estradiol and cultured at a density of $5 \times 10^5$ cells/ml in medium without β-estradiol for 7 days. On day 5, cultures were fed with fresh medium as described above.

For CpG stimulation, 10 nmol of CpG2216 (CpG-A; sequence GGGGGACGATCGTCGGGGGG, Tib Molbiol) or CpG1668 (CpG-B; sequence TCCATGACGTTCCTGATGCT; Tib Molbiol) was complexed with 30 µg DOTAP (Roche/Merck) in HBSS (Gibco/Thermo Scientific) and added at the concentration and for the duration indicated for each experiment. For IFNα stimulation, cells were treated with 100 U/ml IFNα4.

**Tissue preparation, flow cytometry and cell sorting**

BM cells were obtained from mouse femurs and tibias by flushing or centrifugation as previously described (8). Spleens were minced and digested for 30 min at 37°C in PBS containing 1 mg/ml collagenase (Sigma-Aldrich/Merck) and 30 U/ml DNase (Roche/Merck). After washing, RBC lysis, suspensions were passed through a 70-µm cell strainer, resuspended in FACS buffer (PBS, 2 mM EDTA and 2% FBS), and stained. Cultured BM-derived and Hoxb8-FL cells were harvested, washed, and stained. Antibody panels and clone information are provided in Table S8.

Fc receptors were blocked with anti-CD16/CD32 (clone 93) before and during staining. DAPI, 7-AAD, or eFluor 780 was used to identify dead cells. Unless otherwise specified in the relevant figure legend, pDCs were gated as live, single, CD3ε$^-$, CD19$^-$, CD11c$^+$, CD11b$^{-/low}$, B220$^+$, CD317$^+$ and SiglecH$^+$ or as described in the respective figure legends. Flow cytometry was performed using FACSCanto II (BD Biosciences) or FACSFortessa (BD Biosciences) instruments with FACSDiva software, and data were analyzed using FlowJo 10 (BD Biosciences). Cell sorting was performed on a FACSAria III (BD Biosciences) as described previously (8).

**LCMV infection**

LCMV strain WE was provided by Fritz Lehmann-Grube (Heinrich Pette Institute for Immunology and Virology, Hamburg University) and propagated in L929 cells as previously described (9). Mice received 200 PFU of LCMV-WE intravenously. LCMV titers in tissue homogenates were determined using the previously published method (10).

For in vitro infection, MC57 fibroblasts were seeded one day before infection to reach approximately 70% confluence. FACS-purified WT or *Batf*$^{-/-}$ pDCs from BM-derived Flt3L cultures (50,000 cells) were added 18 h before infection. Cultures were infected with LCMV-WE at a multiplicity of infection (MOI) of 0.1 for 4 h. Infection was assessed by staining with the VL-4 rat anti-LCMV monoclonal antibody according to the previously published protocol (10).

**Table S8. Flow cytometry antibody panels**

| Panel | Marker | Fluorophore | Clone/supplier |
|---|---|---|---|
| Ex vivo BATF | CD3 | BV510 | 145-2C11; BD Biosciences |
| Ex vivo BATF | CD19 | BV786 | 1D3; BD Biosciences |
| Ex vivo BATF | CD11c | PE-Cy7 | N418; BioLegend |
| Ex vivo BATF | CD317 / mPDCA1 | BV650 | 927; BioLegend |
| Ex vivo BATF | CD11b | APC | M1/70; BD Biosciences |
| Ex vivo BATF | B220 / CD45R | FITC | RA3-6B2; BD Biosciences |
| Ex vivo BATF | BATF | PE | D7C5; Cell Signaling Technology |
| Ex vivo BATF | Viability dye | eFluor 780 | eBioscience / Thermo Fisher |
| Ex vivo BATF alternative | BATF | Alexa Fluor 647 | D7C5; Cell Signaling Technology |
| Ex vivo BATF alternative | CD11c | APC-Cy7 | N418; BioLegend |
| Ex vivo BATF alternative | CD317 / PDCA1 | PE | 927; eBioscience |
| Ex vivo BATF alternative | CD3ε | PerCP-Cy5.5 | 145-2C11; BD Biosciences |
| Ex vivo BATF alternative | CD19 | PerCP-Cy5.5 | 1D3; BD Biosciences |
| Ex vivo BATF alternative | CD199 / CCR9 | PE-Cy7 | CW-1.2; eBioscience |
| In vitro BATF | CD11b | APC | M1/70; BD Biosciences |
| In vitro BATF | TCRβ | APC-Cy7 | H57-597; BioLegend |
| In vitro BATF | CD19 | APC-Cy7 | 6D5; BioLegend |
| In vitro BATF | SiglecH | BV421 | 440c; BD Biosciences |
| In vitro BATF | CD317 / PDCA1 | BV650 | 927; BioLegend |
| In vitro BATF | BATF | PE | D7C5; Cell Signaling Technology |
| In vitro BATF | CD86 | PerCP-Cy5.5 | GL-1; BioLegend |
| In vitro BATF | CD11c | PE-Cy7 | N418; BioLegend |
| In vitro BATF | B220 / CD45R | FITC | RA3-6B2; BD Biosciences |
| In vitro BATF | Viability dye | eFluor 780 | eBioscience / Thermo Fisher |
| BM pDC sorting | SiglecH | APC | 551; BioLegend |
| BM pDC sorting | CD11b | APC-Cy7 | M1/70; BD Biosciences |
| BM pDC sorting | CD317 / PDCA1 | PE | 927; eBioscience |
| BM pDC sorting | CD3ε | PerCP-Cy5.5 | 145-2C11; BD Biosciences |
| BM pDC sorting | CD19 | PerCP-Cy5.5 | 1D3; BD Biosciences |
| BM pDC sorting | CD11c | PE-Cy7 | N418; BioLegend |
| BM pDC sorting | B220 / CD45R | FITC | RA3-6B2; BD Biosciences |
| BM pDC sorting | Viability dye | 7-AAD | BD Bioscience |
| Chimera BM pDC sorting | B220 / CD45R | APC | RA3-6B2; BD Biosciences |
| Chimera BM pDC sorting | CD11b | APC-Cy7 | M1/70; BD Biosciences |
| Chimera BM pDC sorting | CD317 / PDCA1 | PE | 927; eBioscience |
| Chimera BM pDC sorting | CD3ε | PerCP-Cy5.5 | 145-2C11; BD Biosciences |
| Chimera BM pDC sorting | CD19 | PerCP-Cy5.5 | 1D3; BD Biosciences |
| Chimera BM pDC sorting | CD11c | PE-Cy7 | N418; BioLegend |
| Chimera BM pDC sorting | CD45.2 | FITC | 104; BD Biosciences |
| Chimera BM pDC sorting | Viability dye | 7-AAD | BD Biosciences |
| All panels | Fc receptor block, CD16/CD32 | Unconjugated | 93; eBioscience |

### RT-qPCR

Total RNA was isolated from FACS-purified resting or CpG-stimulated pDCs or homogenized tissues using the NucleoSpin RNA Mini Kit (Macherey-Nagel). RNA concentration and quality were assessed using a NanoDrop 1000 spectrophotometer (PEQLab). cDNA was synthesized with SuperScript III reverse transcriptase and oligo(dT) (Thermo Scientific) according to the manufacturer's instructions. Amplifications were performed on a CFX96 Real-Time C1000 Thermal Cycler (Bio-Rad) using FastStar TaqMan Probe Master Mix from Roche for *Ifnb1*, *Batf*, *Batf2*, *Zfp366*, and the respective probe library (Roche). Transcripts encoding IFNα were measured using wobble primers targeting several *Ifna* subtypes. *Ifna* and *Batf3* transcripts were quantified using MESA GREEN qPCR Master Mix Plus for SYBR Assay (Eurogentec). *Actb* (β-actin) was used as a reference housekeeping gene for quantification. Table S2 lists the primer sequences and probe identifiers used for RT-qPCR.

### Cytokine ELISA and multiplex profiling

FACS-purified pDCs were seeded at 1 × $10^5$ cells/well in 200 µl in 96-well plates or 5 × $10^5$ cells/well in 500 µl in 24-well plates. After at least 1 h at 37°C, cells were stimulated with the indicated concentrations of CpG2216 or CpG1668 complexed with DOTAP. IFNα, IFNβ and IL-12p40 were measured in cell-free supernatants or mouse serum using mouse IFNα ELISA (Invitrogen/Thermo Scientific), IFNβ ELISA (BioLegend), and IL-12p40 ELISA (R&D Systems), respectively. For multiplex analysis, purified pDCs were stimulated for 16 h, and 36 cytokines/chemokines were measured using a ProcartaPlex mouse Cytokine/Chemokine Panel and a Luminex 100 instrument according to the manufacturer's instructions.

**Table S9. RT-qPCR primers and probes**

| **Target** | Forward primer | *Reverse primer* | Probe/chemistry |
|---|---|---|---|
| *Ifnb1* | CCT TTG ACC TTT CAA ATG CAG | *CAG GCA ACC TTT AAG CAT CAG* | Universal ProbeLibrary #95 |
| *Ifna* (wobble) | ATG GCT AGR CTC TGT GCT TTC CT | *AGG GCT CTC CAG AYT TCT GCT CTG* | SYBR Green |
| *Batf* | AGA AAG CCG ACA CCC TTC | *CGG AGA GCT GCG TTC TGT* | Universal ProbeLibrary #85 |
| *Batf2* | GAA GCA GAA GAA CCG AGT GG | *TCT CCA AGG ATT CGT GCT G* | Universal ProbeLibrary #29 |
| *Batf3* | CAG AGC CCC AAG GAC GAT G | *GCA CAA AGT TCA TAG GAC ACA GC* | SYBR Green |
| *Zfp366* | AGG CCT GCA CTC TCA AGG T | *TCT GGG AGA GGT CAA ATG GT* | Universal ProbeLibrary #15 |
| *Actb* | CGC TCA GGA GGA GCA ATG | *TGA CAG GAT GCA GAA GGA GA* | Universal ProbeLibrary #106 |
| *Isg15* | AGT CGA CCC AGT CTC TGA CTC T | *CCC CAG CAT CTT CAC CTT TA* | Universal ProbeLibrary #71 |
| *Isg20* | GCG CCT GCT ACA CAA GAA C | *CTG AGA GAT TTT GTA GAG CTC CAT T* | Universal ProbeLibrary #81 |
| *Mx1* | TTC AAG GAT CAC TCA TAC TTC AGC | *GGG AGG TGA GCT CCT CAG T* | Universal ProbeLibrary #53 |

### Immunoprecipitation and immunoblotting

For immunoprecipitation, proteins were transiently expressed using JetPRIME (Polypus). After 2 days, cells were washed twice with PBS and lysed for 30 min at 4°C in modified RIPA buffer (50 mM Tris-HCl, 150 mM NaCl, 1% NP-40, 0.5% sodium deoxycholate, 1 mM Na3VO5, 1 mM EDTA, 1 mM PMSF, and protease inhibitor cocktail), followed by three freeze-thaw cycles. Lysates were clarified at 20,000 × g for 20 min at 4°C and incubated overnight with EZview Red anti-c-Myc affinity gel (Merck). Beads were washed five times with lysis buffer; bound proteins were eluted in Laemmli buffer, separated on NuPAGE 4-12% Bis-Tris gels, and transferred to nitrocellulose membranes (GE Healthcare Life Sciences). Membranes were probed with anti-IRF7 (polyclonal antibody, Merck Millipore, ABF130), anti-Myc (clone 9E10, Merck Millipore, MABE282), anti-BATF (clone D7C5, Cell Signaling Technology, 8638), or anti-GAPDH (polyclonal, Abcam, ab9485), as appropriate. HRP-conjugated secondary antibodies (goat anti-mouse [Jackson ImmunoResearch Laboratories, 15-035-068] or mouse anti-rabbit [Jackson ImmunoResearch Laboratories, 211-032-171]) and a chemiluminescent substrate (SuperSignal chemiluminescent substrate; Thermo Scientific) were used for detection. For whole-cell lysates, 10 µg protein was loaded per sample unless otherwise indicated.

### Plasmids and dual-luciferase reporter assays

pmBatf, pmSpi1 and pZfp366-Myc-DDK encode murine BATF, PU.1 and DC-SCRIPT, respectively. All three expression vectors were purchased from OriGene Technologies GmbH. pRL-CMV (Promega) encodes Renilla luciferase under the CMV promoter. The pIFNβ-Luc and pIFNα4-Luc reporter plasmids, based on pGL3basic (Promega), contain firefly luciferase under the *Ifnb1* and *Ifna4* promoters, respectively. The *Zfp366* promoter reporters pGL4_Zfp366P-3904, -2900, -2173 and -1202 contain promoter fragments extending from 100 bp downstream of the transcription start site to the indicated upstream positions. To generate these reporter plasmids, PCR-amplified *Zfp366* promoter sequences were inserted into the pGL4.17_luc2/Neo plasmid (Promega), upstream of the firefly luciferase coding sequence using In-Fusion Snap Assembly Master Mix (Takara Bio). pBEC2VmIRF7a and pFlag-IRF-7 express murine and human IRF7, respectively; phTBK1-Flag-His expresses human TBK1. All these plasmids were kind gifts from Prof. Dr. Stefan Bauer (Philipps University Marburg, Germany).

pWPI is a bicistronic lentiviral vector in which an EGFP-encoding sequence is inserted downstream of an EMCV IRES site. The plasmid expresses EGFP under the control of the constitutive EF1α promoter and was a kind gift from Prof. Didier Trono (Addgene plasmid #12254). pWPI-mBatf-GFP was generated by ligating a murine BATF-encoding sequence into the pWPI expression plasmid using the SwaI and PacI restriction sites upstream of the IRES sequence. pWPI-mZfp366-GFP encodes murine DC-SCRIPT and EGFP. PCR-amplified murine *Zfp366* coding sequences were inserted at the SwaI and PacI sites into the pWPI expression plasmid to construct the pWPI-mZfp366-GFP bicistronic expression plasmid. The lentiviral envelope vector pLP/VSVG was purchased from Invitrogen (Life Technologies), and the packaging vector psPAX2 was a kind gift from Prof. Didier Trono (Addgene plasmid #12260).

The activity of the *Ifnb1* promoter or the *Ifna4* promoter was analyzed using an IRF7-dependent dual-luciferase reporter assay as previously described (11, 12). Briefly, for *Ifnb1* and *Ifna4* reporter assays, $1.25 \times 10^5$ 293FT cells were seeded per well in 24-well plates. The following day, cells were co-transfected with 90 ng of the firefly reporter, 10 ng of pRL-CMV, 10 ng of the IRF7 expression vector, 40 ng of the TBK1 expression vector, and the indicated amounts of BATF-, DC-SCRIPT- or control vector. For *Zfp366* promoter assays, cells received 90 ng of the reporter, 10 ng of pRL-CMV, 10 ng of the PU.1 vector where indicated, and the indicated amounts of BATF or control vector. Total DNA was kept constant using an empty vector. Transfection was performed using polyethylenimine (PEI; Sigma-Aldrich/Merck) (13, 14). After 20 h, cells were lysed in passive lysis buffer (Promega) and luciferase activity was measured using a dual-luciferase kit (Promega), and a Mithras LB 940 luminometer (Berthold Technologies). Two seconds after the addition of 50 µl of substrate solution to 10 µl of cell lysate, firefly luciferase activity was measured for 10 seconds, followed by measurement of Renilla luciferase activity for 5 seconds. Firefly activity was normalized to Renilla activity.

**RNA-seq**

One million FACS-purified BM-derived Flt3L-cultured pDCs were left untreated or stimulated with 1 µM CpG2216 complexed with DOTAP for 2, 6 or 12 h. Total RNA was isolated using the NucleoSpin RNA Mini Kit (Macherey-Nagel), DNase-treated and quantified using the Qubit RNA HS Assay (Thermo Fisher Scientific). RNA quality was assessed by capillary electrophoresis using a Fragment Analyzer and the ‘Total RNA Standard Sensitivity Assay’ (Agilent Technologies). All samples in this study were of high quality, with a mean RNA quality number of 9.9.

Libraries were prepared from 200 ng of total RNA using the Illumina TruSeq Stranded mRNA Library Prep Kit following the manufacturer’s protocol. The bead-purified libraries were subsequently normalized and sequenced on a HiSeq 3000/4000 system (Illumina, Inc.) as single-end 150-bp reads. The bcl2fastq tool was used to convert the BCL files to FASTQ files as well as for adapter trimming and demultiplexing.

Reads were processed in CLC Genomics Workbench v11.0.1 (Qiagen), adapter-trimmed (Illumina TruSeq) and quality-trimmed (using the default parameters, bases below Q13; a maximum of two ambiguous nucleotides), and mapped to the mouse mm10 (GRCm38.86) reference genome. Three biological replicates were included per condition. Samples were grouped according to their respective experimental conditions. Raw counts were subsequently uploaded to the Galaxy web platform. Multigroup comparisons were performed using the public server at usegalaxy.org (15). Pearson correlations were used for replicate quality control. Our results revealed high similarity (greater than 95%) among the biological replicates within the respective conditions of the RNA-seq dataset. Differential gene expression between any two conditions was calculated using the edgeR quasi-likelihood pipeline, which uses negative binomial generalized linear models with the F test (16, 17). Lowly expressed genes were filtered using a counts-per-million (CPM) threshold, which was calculated based on the average library size of our NGS run (18). P values were adjusted using false discovery rate (FDR) correction; adjusted P ≤ 0.05 was considered significant.

Publicly available human data (GSE93679) were analyzed using the Biobase, GEOquery and limma R packages (19, 20).

**BATF ChIP-seq**

FACS-purified BM-derived Flt3L-cultured pDCs from C57BL/6N mice were left untreated or stimulated for 2 h with 1 µM CpG2216 complexed with DOTAP. Cells were fixed in 1% formaldehyde for 15 min and quenched with 0.125 M glycine. ChIP-seq was performed by Active Motif Services (Carlsbad, California, USA). Chromatin was isolated by the addition of lysis buffer, followed by disruption with a Dounce homogenizer. The lysates were sonicated and the DNA was sheared to an average length of 300-500 bp. Genomic DNA (input) was prepared by treating aliquots of chromatin with RNase, proteinase K and heat for decrosslinking, followed by ethanol precipitation. The pellets were resuspended, and the resulting DNA was quantified on a NanoDrop spectrophotometer. Extrapolation to the original chromatin volume allowed quantitation of the total chromatin yield. For immunoprecipitation, 20 µg of chromatin was spiked with 200 ng of Drosophila chromatin and precleared with protein A agarose beads. BATF was immunoprecipitated using 4 µg of anti-BATF antibody (clone D7C5; Cell Signaling Technology, 8638BF); 0.4 µg of anti-H2Av antibody was included for spike-in chromatin recovery (21). The complexes were washed, eluted from the beads with SDS buffer, and subjected to RNase and proteinase K treatment. Crosslinks were reversed by incubation overnight at 65°C, and the ChIP DNA was purified via phenol-chloroform extraction and ethanol precipitation. Quantitative PCR (qPCR) reactions were carried out in triplicate on specific genomic regions using SYBR Green Supermix (Bio-Rad). The resulting signals were normalized for primer efficiency by carrying out qPCR for each primer pair using input DNA.

For ChIP sequencing, Illumina sequencing libraries were prepared from the ChIP and input DNAs by the standard consecutive enzymatic steps of end-polishing, dA addition, and adaptor ligation. The steps were performed on an automated system (Apollo 342, Wafergen Biosystems/Takara). After a final PCR amplification step, the resulting DNA libraries were quantified and sequenced on an Illumina NextSeq 500 (75 nt reads, single end). Reads were aligned consecutively to the mouse genome (mm10) and to the Drosophila genome (dm3) using the BWA algorithm (default settings). Duplicate reads were removed, and only uniquely mapped reads (mapping quality ≥ 25) were used for further analysis. The number of mouse alignments used in the analysis was adjusted according to the number of Drosophila alignments that were counted in the samples being compared. Mouse alignments were extended in silico at their 3′ ends to a length of 200 bp, which is the average genomic fragment length in the size-selected library, and assigned to 32-nt bins along the genome. The resulting histograms (genomic “signal maps”) were stored in bigWig files. Peak locations were determined using the MACS algorithm (v2.1.0) with a cutoff of P = 1e-7. Peaks that were on the ENCODE blacklist of known false ChIP-seq peaks were removed. Signal maps and peak locations were used as input data for Active Motifs proprietary analysis program, which creates Excel tables containing detailed information on sample comparison, peak metrics, peak locations, and gene annotations. Using BATF ChIP-seq, a qualitative distribution of ChIP-seq peaks was determined based on their genomic location (such as introns, 3’-UTRs, distal (1-3 kb) and proximal (0-1 kb) promoter regions). Of the 15,453 BATF ChIP-seq peaks, 4,470 and 10,983 were located within genes annotated by RefSeq in untreated and CpG-stimulated pDCs, respectively. The results were further visualized using Integrative Genomics Viewer (IGV) (22) and modified with Inkscape or Adobe Illustrator. To present ChIP-seq average peak values (Avg Peak), the normalized average signal intensity across all peaks associated with a given gene was calculated by dividing the sum of all peak signals by the total number of peaks assigned to that gene. For example, if a gene has multiple peaks (e.g., 8), the individual peak values are combined, and an average is computed to provide a single representative measurement.

#### ATAC-seq

Viable FACS-purified BM-derived Flt3L-cultured pDCs from WT (*Batf*$^{+/+}$) and Batf$^{-/-}$ mice were left untreated or stimulated with 1 µM CpG2216 for 2 h. Cells were harvested and snap-frozen in culture medium supplemented with FBS and 5% DMSO before shipment to Active Motif Services (Carlsbad, California, USA). After thawing, cells were washed in cold PBS, lysed and subjected to transposase-mediated tagmentation as previously described (23), with modifications based on Corces *et al.* (24). Briefly, cell pellets were resuspended in lysis buffer, pelleted, and tagmented using the enzyme and buffer provided in the Nextera Library Prep Kit (Illumina). The tagmented DNA was purified using a MinElute PCR Purification Kit (Qiagen), amplified for 10 PCR cycles and purified using Agencourt AMPure SPRI beads (Beckman Coulter). The resulting material was quantified using the KAPA Library Quantification Kit for Illumina platforms (KAPA Biosystems) and sequenced on a NextSeq 500 sequencer (Illumina) using paired-end 42-bp reads.

Analysis of ATAC-seq data was very similar to the analysis of ChIP-seq data. Reads were aligned to mm10 using the BWA algorithm (MEM mode; default settings). Duplicate reads were removed; matched pairs with mapping quality ≥1 were retained. Alignments were extended in silico at their 3′ ends to 200 bp and assigned to 32-nt genomic bins. The resulting histograms (genomic “signal maps”) were stored in bigWig files. Peaks were called with the MACS2 v2.1.0 algorithm at a cutoff p value of 1e-7, without a control file, and with the –nomodel option. Peaks that were on the ENCODE blacklist of known false ChIP-seq peaks were removed. Signal maps and peak locations were used as input data to Active Motifs proprietary analysis program, which creates Excel tables containing detailed information on sample comparisons, peak metrics, peak locations, and gene annotations. Pearson correlation analysis for the ATAC-seq data showed > 95% similarity among all the biological replicates, indicating a high data quality. For differential analysis, reads were counted in merged peak regions using Subread, and differential accessibility was analyzed using DESeq2.

#### Omics visualization, human gene expression and genetic analyses

Volcano plots and MA plots were generated using ggplot2 (25) and ggrepel (26). Heatmaps were generated using the Morpheus online tool (Broad Institute; https://software.broadinstitute.org/morpheus). Venn diagrams were created using eulerr (27). Pathway analyses were performed using the Reactome database (28). GSEA was performed using GSEA software (29) and Hallmark gene sets (30). Network visualization of the omics data was performed in Cytoscape (31). ChIP-seq tracks and peaks were visualized using Integrative Genomics Viewer.

The publicly available human expression dataset GSE93679 was analyzed using Biobase, GEOquery and limma. Genetic-association data for the *BATF* and *ZNF366* loci were retrieved from the GWAS Catalog (32) (accessed 18 June 2026; Table S7) and visualized as regional association plots generated in R (version 4.5.2) using RStudio (version 2026.01.0, build 392) and ggplot2 (version 4.0.3). Variants are plotted at their GRCh38 genomic positions, with one entry per variant representing the most significant reported association. Variants of interest are highlighted in yellow. Gene positions and strands orientations were obtained from Ensembl release 116 (GRCh38). Where necessary, genomic coordinates were converted from GRCh38 to hg19 genome coordinates using the Lift Genome Annotations tool in the UCSC Genome Browser (https://genome.ucsc.edu/index.html) (33). AnimalTFDB was used to classify transcription factors, cofactors and chromatin-remodeling factors. AnimalTFDB was previously generated by systematically scanning animal proteomes for signature

DNA-binding and regulatory domains, classifying proteins into transcription factors, transcription cofactors, and chromatin-remodeling factors, annotating them with functional annotations, homology data, and manual curation (34). The database provides genome-wide coverage of transcription-related proteins (transcription factors, co-factors, and chromatin-remodeling factors) across 183 animal genomes (35) and is widely used for categorizing transcription-related proteins, providing TF family assignments alongside TF-binding-site information based on peer-reviewed publications.

#### Quantification and statistical analysis

Statistical analyses were performed using GraphPad Prism 9. Student's t test or one-way or two-way ANOVA with multiple-comparison post-tests was used as specified in the figure legends. $P < 0.05$ was considered statistically significant. Exact n values, the definition of biological or technical replicates, and the specific post-test are stated in the relevant figure legends.

#### Supporting Information: References

15. E. Afgan *et al.*, The Galaxy platform for accessible, reproducible and collaborative biomedical analyses: 2016 update. *Nucleic Acids Res* **44**, W3–W10 (2016).
16. M. D. Robinson, D. J. McCarthy, G. K. Smyth, edgeR: a Bioconductor package for differential expression analysis of digital gene expression data. *Bioinformatics* **26**, 139–140 (2010).
17. R. Liu *et al.*, Why weight? Modelling sample and observational level variability improves power in RNA-seq analyses. *Nucleic Acids Res* **43**, e97 (2015).
18. Y. Chen, A. T. Lun, G. K. Smyth, From reads to genes to pathways: differential expression analysis of RNA-Seq experiments using Rsubread and the edgeR quasi-likelihood pipeline. *F1000Res* **5**, 1438 (2016).
19. S. Davis, P. S. Meltzer, GEOquery: a bridge between the Gene Expression Omnibus (GEO) and BioConductor. *Bioinformatics* **23**, 1846–1847 (2007).
20. G. K. Smyth, Linear models and empirical bayes methods for assessing differential expression in microarray experiments. *Stat Appl Genet Mol Biol* **3**, Article3 (2004).
21. B. Egan *et al.*, An Alternative Approach to ChIP-Seq Normalization Enables Detection of Genome-Wide Changes in Histone H3 Lysine 27 Trimethylation upon EZH2 Inhibition. *PLoS One* **11**, e0166438 (2016).
22. J. T. Robinson *et al.*, Integrative genomics viewer. *Nat Biotechnol* **29**, 24–26 (2011).
23. J. D. Buenrostro, P. G. Giresi, L. C. Zaba, H. Y. Chang, W. J. Greenleaf, Transposition of native chromatin for fast and sensitive epigenomic profiling of open chromatin, DNA-binding proteins and nucleosome position. *Nat Methods* **10**, 1213–1218 (2013).
24. M. R. Corces *et al.*, An improved ATAC-seq protocol reduces background and enables interrogation of frozen tissues. *Nat Methods* **14**, 959–962 (2017).
25. H. Wickham, *ggplot2: Elegant Graphics for Data Analysis* (Springer-Verlag, New York, 2016).
26. K. Slowikowski (2020) ggrepel: Automatically Position Non-Overlapping Text Labels with 'ggplot2.
27. J. Larsson (2018) eulerr: Area-Proportional Euler and Venn Diagrams with Ellipses.
28. B. Jassal *et al.*, The reactome pathway knowledgebase. *Nucleic Acids Res* **48**, D498–D503 (2020).
29. A. Subramanian *et al.*, Gene set enrichment analysis: a knowledge-based approach for interpreting genome-wide expression profiles. *Proc Natl Acad Sci U S A* **102**, 15545–15550 (2005).
30. A. Liberzon *et al.*, The Molecular Signatures Database (MSigDB) hallmark gene set collection. *Cell Syst* **1**, 417–425 (2015).
31. P. Shannon *et al.*, Cytoscape: a software environment for integrated models of biomolecular interaction networks. *Genome Res* **13**, 2498–2504 (2003).
32. A. Buniello *et al.*, The NHGRI-EBI GWAS Catalog of published genome-wide association studies, targeted arrays and summary statistics 2019. *Nucleic Acids Res* **47**, D1005–D1012 (2019).
33. D. Karolchik *et al.*, The UCSC Genome Browser database: 2014 update. *Nucleic Acids Res* **42**, D764–770 (2014).
34. H. M. Zhang *et al.*, AnimalTFDB: a comprehensive animal transcription factor database. *Nucleic Acids Res* **40**, D144–149 (2012).
35. W. K. Shen *et al.*, AnimalTFDB 4.0: a comprehensive animal transcription factor database updated with variation and expression annotations. *Nucleic Acids Res* **51**, D39–D45 (2023).
